# Tumor nutrient stress gives rise to a drug tolerant cell state in pancreatic cancer

**DOI:** 10.1101/2025.09.04.673818

**Authors:** Colin Sheehan, Lyndon Hu, Guillaume Cognet, Grace Croley, Thao Trang Nguyen, Anika Thomas-Toth, Darby Agovino, Patrick B. Jonker, Mumina Sadullozoda, Leah M. Ziolkowski, James K. Martin, Alica K. Michels, Ranya Dano, Mohammed A. Khan, Rasangi Perera, Isabel N. Alcazar, Christopher J. Halbrook, Kay F. Macleod, Hardik Shah, Christopher R. Weber, James L. LaBelle, Alexander Muir

## Abstract

Systemic therapies are the standard of care for most pancreatic ductal adenocarcinoma (PDAC) patients but provide limited benefit due to pervasive resistance. The fibrotic tumor microenvironment (TME) is thought to drive resistance by restricting perfusion and drug delivery. Here, we show that therapeutically relevant drug concentrations are achieved even in poorly perfused, therapy-resistant murine PDAC tumors, indicating that impaired delivery alone does not explain drug resistance. Instead, we find TME exposure imprints a therapy-resistant state upon PDAC cells. These observations raised the question of how the TME imposes this state. Poor perfusion alters nutrient availability in the TME. To model this, we developed Tumor Interstitial Fluid Medium (TIFM), which recapitulates TME nutrient conditions. TIFM cultured PDAC cells acquire a therapy-resistant phenotype that mirrors resistance observed in the TME. In this state, cytotoxic and targeted therapies retain on-target activity but fail to trigger cell death, resulting in therapeutic tolerance. Mechanistically, drug tolerance is driven by suppression of apoptotic priming and can be reversed by inhibition of the anti-apoptotic regulator BCL-XL. These results identify TME-driven reprogramming of cell death as a key mechanism of therapy resistance in PDAC and establish TIFM as a physiologically relevant model for studying microenvironment-induced drug resistance.

## Introduction

Pancreatic ductal adenocarcinoma (PDAC) remains one of the deadliest malignancies with a 5-year survival rate of only 12% (1). This low survival rate is driven by multiple factors including the inability to surgically resect most PDAC tumors and chemoresistance exhibited by non-resected tumors (2). Indeed, the majority of PDAC patients exhibit primary resistance to aggressive combination chemotherapy regimens such as gemcitabine and nab-paclitaxel or FOLFIRINOX (fluorouracil, irinotecan and oxaliplatin) (3, 4). Only the minority of patients display objective tumor responses with complete responses rarely reported (3, 4). More recently, KRAS inhibitors, which target the most common oncogenic driver in PDAC, have shown efficacy in the treatment of murine (5–7) and human (8) PDAC. However, even for these targeted therapies, resistance commonly occurs (9). Thus, the inability to reliably control PDAC with both targeted and chemotherapies remains a significant barrier to effective treatment in this disease.

Understanding the biological processes underlying drug resistance in PDAC could enable new approaches to improve therapeutic efficacy in this disease. Studies from human and mouse PDAC point to the tumor microenvironment (TME) as playing a critical role in drug resistance (10). While patients are largely unresponsive to chemotherapies (3, 4), cultured human PDAC cell lines are eradicated at very low concentrations of the same drugs (11). Similarly, both genetically engineered and orthotopic allograft mouse models of PDAC are highly resistant to chemotherapeutics (12, 13), but cell lines established from these models respond to low doses of these drugs (13). Thus, both human and murine PDAC cells are not inherently therapy resistant. Rather, drug resistance must be imparted by the TME.

The PDAC TME has been proposed to inhibit chemotherapy response by impairing the ability of drugs to reach PDAC cells. PDAC tumors are both hypovascularized (14) and exhibit dramatic desmoplasia (15) where the stroma can compress and compromise the functionality of blood vessels in the tumor (12, 16, 17). Together, these properties of PDAC substantially limit perfusion in the tumor (18). Limited perfusion correlates with poor patient outcomes (18) and therapy resistance (19), which has led to a search to understand and overcome the mechanisms by which limited perfusion in the PDAC TME drives therapy resistance. One mechanism by which poor perfusion is thought to drive therapy resistance is by limiting the delivery of therapeutic agents into the TME (20). Indeed, drug delivery to PDAC tumors can be impaired by the TME (12, 16, 17). However, recent preclinical and clinical studies have shown that approaches to improve drug penetrance in PDAC tumors, such as direct delivery of therapeutic agents to the tumor parenchyma (21) or the use of agents targeting the PDAC desmoplastic reaction (22–25), do not improve outcomes. Together, these results argue for the existence of additional mechanisms by which poor perfusion in the TME causes chemoresistance, beyond being a barrier to drug delivery (26).

In this study, we sought to understand how limited perfusion contributes to therapy resistance in PDAC. Using a murine model of PDAC that recapitulates both the poor perfusion and drug resistance observed in human PDAC, we first examined the extent to which impaired drug delivery contributes to drug resistance. We found that despite perfusion deficits in these PDAC tumors, drugs accumulate to therapeutic levels in the TME, and treated tumors exhibit evidence of on-target drug activity. These findings suggest that poor perfusion limits drug efficacy by mechanisms beyond impaired drug delivery. One consequence of poor perfusion is abnormal nutrient availability and metabolic stress in the TME (27). To study the contribution of TME metabolic stress as a contributor to therapy resistance in poorly perfused PDAC, we cultured PDAC cells in Tumor Interstitial Fluid Medium (TIFM), a culture medium that recapitulates the nutrient availability of the PDAC TME (28). TIFM culture resulted in PDAC cells acquiring resistance to both targeted and chemotherapies, which was driven not driven by suppression of drug activity, but rather by tolerance of PDAC cells to cytotoxic stress induced by therapeutics. This drug tolerant phenotype was enabled by suppression of apoptotic priming in PDAC cells and could be overcome by targeting the anti-apoptotic regulator BCL-XL. Consistent with these findings, PDAC tumors contain drug tolerant cells with suppressed apoptotic priming. Taken together, these studies indicate that poor perfusion in PDAC does not causes drug resistance by preventing access of drugs to the TME. Rather, pathophysiological nutrient levels that occur in the poorly perfused TME imprint a state of suppressed apoptotic priming that enables PDAC cells to tolerate drug-induced cytotoxic stress.

## Results

### Poor perfusion is associated with reduced therapy response in PDAC

Given previous work correlating PDAC perfusion rates with therapy response and outcomes (18, 19), we sought to understand how perfusion deficits in PDAC influence drug sensitivity. To do so, we used an orthotopic allograft model of PDAC (28), which we found did not significantly respond to high dose treatment with the clinically-used chemotherapeutic agent, gemcitabine (Figure 1A,B), mirroring drug resistance observed in human PDAC and other animal models of this disease. We then measured perfusion these tumors by monitoring the delivery of the perfusion tracer Hoechst 33342 from the vascular space to the tumor parenchyma (Figure 1C). Hoechst 33342 is a membrane-permeable, DNA-binding fluorogenic dye, with extremely short half-life in the circulation (<2 minutes), enabling labeling of perfused cells within tissues (29). Hoechst 33342 labeling indicated that PDAC tumors were far less perfused than healthy pancreata (Figure 1D,E) or tumor-adjacent pancreas (Figure 1F,G). Interestingly, Hoechst 33342 labeling in tumors exhibited a mottled pattern of labeling unevenly across the tumor (Figure 1D,F). To understand the cause of heterogenous perfusion in PDAC tumors, we used iterative multiplexed immunofluorescence imaging (t-CyCIF) (30), to assess colocalization of Hoechst labeling with the vascular biomarker CD31. Furthermore, to identify functional blood vessels, we intravenously delivered fluorescently conjugated *L. Esculentum* lectin (LEL) to label blood vessels actively transporting blood (Figure 1H) as previously described (12). We found that Hoechst 33342 labeling in the tumor strongly correlated with proximity to LEL+ CD31+ blood vessels (Figure 1I,J). Interestingly, we found a substantial decrease of LEL labeling in CD31+ vessels in PDAC tumors compared to normal pancreata (Figure 1K). This indicates that many blood vessels are functionally impaired in our animal model of PDAC, as has been observed in other animal models of PDAC (12). The observation that LEL-vessels did not support robust Hoechst 33342 labeling of the surrounding tumor (Figure 1I) is consistent with prior findings that stromal compression and impairment of tumor vasculature can affect PDAC perfusion (12, 16, 17). Collectively, these results demonstrate that impairments in blood vessel function lead to impaired perfusion in PDAC tumors.

**Figure 1:**
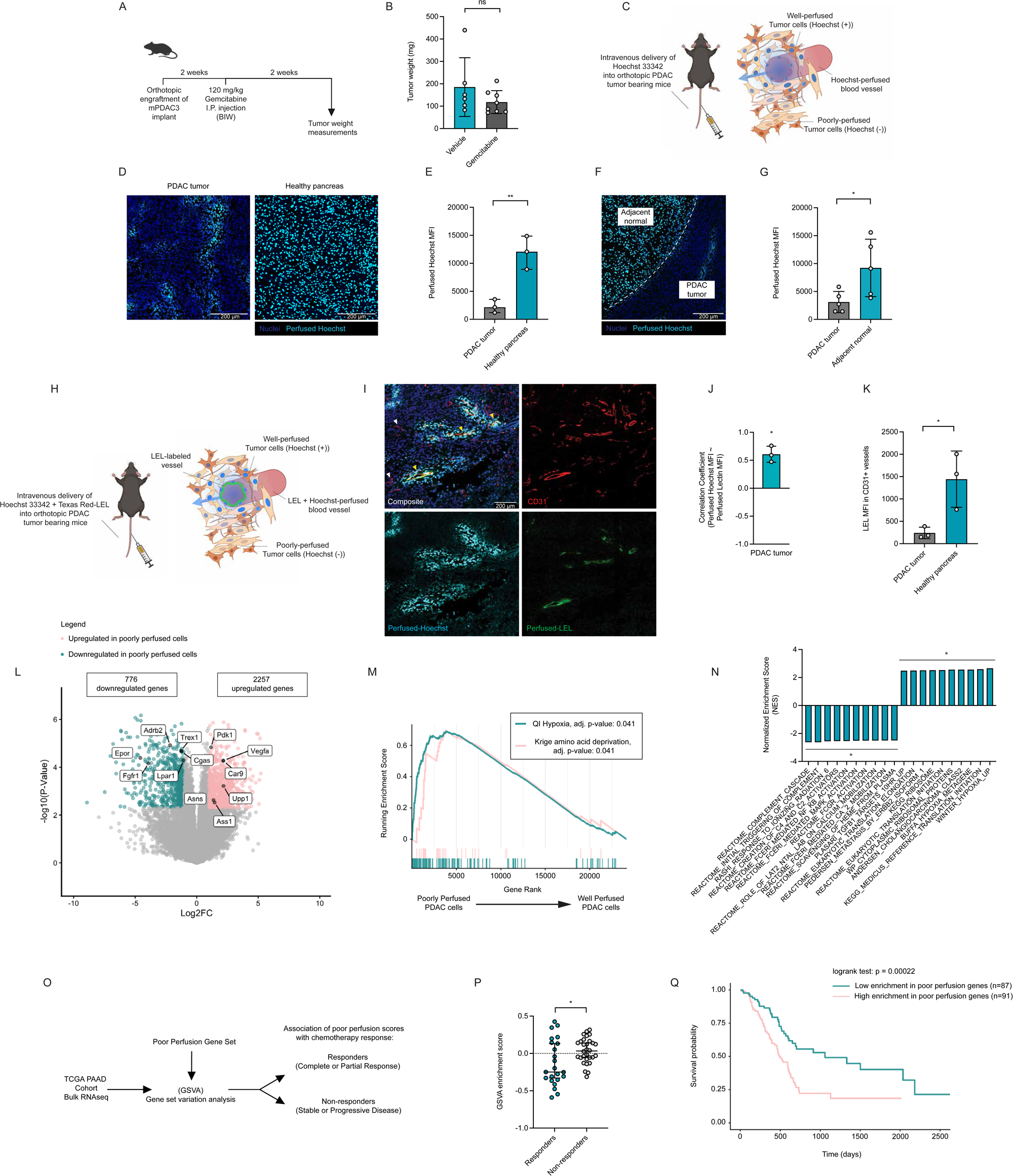
PDAC tumors are poorly perfused and perfusion levels correlate with PDAC therapy response. **(A)** Diagram of workflow for establishing orthotopic PDAC models with mPDAC3-RPMI cells and treatment of tumor bearing animals with gemcitabine. **(B)** mPDAC3-RPMI tumor weights from animals treated with vehicle (n = 6) or gemcitabine (n = 8). **(C)** Diagram of workflow for intravenous injection of mice with Hoechst 33342 to label well-perfused cells in tumor-bearing mice. **(D)** Representative images of Hoechst 33342 staining (cyan) of orthotopic PDAC tumors and normal pancreata. Total nuclei in blue. Scale bars correspond to 200 µm. **(E)** Quantification of perfused Hoechst 33342 labeling (median fluorescence intensity per nuclei) in orthotopic PDAC tumors (n = 3) and normal pancreata (n = 3). **(F)** Representative image of orthotopic PDAC tumor with adjacent normal pancreas perfused with Hoechst 33342 (cyan). Dashed line denotes boundary between tumor and normal tissue. Total nuclei in blue. Scale bar corresponds to 200 µm. **(G)** Quantification of Hoechst 33342 labeling (median fluorescence intensity per nuclei) in PDAC tumors and adjacent normal pancreas (n = 5). **(H)** Diagram of workflow for intravenous injection of mice with Hoechst 33342 and *L. Esculentum* lectin (LEL) to label functional blood vessels and perfused cells in tumor bearing mice. **(I)** Representative images of Hoechst 33342 (cyan), anti-CD31 (red), LEL (green) and nuclei (blue) staining of PDAC tumors. Scale bar corresponds to 200 µm. Yellow arrowheads indicate perfused vessels (LEL+, Hoechst+, CD31+) while white arrowheads indicate non-perfused vessels (LEL-, Hoechst-, CD31+). **(J)** Correlation coefficients of LEL labeling in blood vessels and Hoechst 33342 staining intensity in vessel-adjacent tumor cells in PDAC tumors (n = 3). **(K)** Quantification of median fluorescence intensity of LEL labeling in CD31+ vessels in PDAC tumors (n = 3) and normal pancreata (n = 3). **(L)** Volcano plot of differentially expressed genes (DEGs) between Hoechst 33342-high (well perfused) and Hoechst 33342-low (poorly perfused) mPDAC3-RPMI cells from PDAC tumors (n = 4). Blue: downregulated in poorly perfused PDAC cells p < 0.05. Pink: upregulated genes in poorly perfused PDAC cells with adjusted p < 0.05. Gray: genes with adjusted p > 0.05. Adjusted p-value was calculated using Benjamini-Hotchberg (BH) false discovery rate method. **(M)** Gene set enrichment analysis of “QI Hypoxia” and “Krige amino acid deprivation” gene sets in poor and well perfused PDAC cells. Adjusted p-values calculated following analysis of all MSigDB curated C2 gene sets using Benjamini-Hotchberg (BH) false discovery rate method. **(N)** Top enriched MSigDB curated C2 gene sets in poor and well perfused mPDAC3-RPMI cells. All gene sets displayed have an adjusted p-value < 0.05, determined by Benjamini-Hotchberg (BH) method. **(O)** Diagram for assessing expression of genes associated with poorly perfused PDAC in the TGCA PDAC cohort. **(P)** Gene set variation analysis (GSVA) enrichment scores for poor perfusion marker genes in TGCA PDAC patients who responded (complete or partial responses) or not (stable or progressive disease) to chemotherapeutic treatment. **(Q)** TGCA PDAC patients were stratified into cohorts with high (n = 89) or low (n = 89) GSVA enrichment of poor perfusion marker genes and Kaplan Meier survival analysis was performed. Statistical significance was determined by log-rank test. Statistical significance in B, E, G, K, P was determined using 2 sample T-test. For J, correlation coefficients between perfused lectin MFI and perfused Hoechst 33342 MFI of vessels were calculated by spearman’s correlation (n=1000), and a statistically significant difference from 0 of the collective correlation coefficients was determined by one-sample T-test (n=3). * p≤0.05 ** p≤0.01 *** p≤0.001 and **** p≤0.0001.

We next sought to determine how perfusion level impacts PDAC cell state. We identified malignant cells from PDAC tumors using the PDAC cell surface marker mesothelin (MSLN) (31) and sorted cells that were highly and lowly labeled with Hoechst 33342 (Supplemental Figure 1A). We then characterized the transcriptome of these well and poorly perfused PDAC cells. Perfusion status altered the expression of over 3000 genes in PDAC cells, indicating that perfusion is a major regulator of PDAC gene expression (Figure 1L). At the individual gene level, poorly perfused PDAC cells displayed upregulation of a number of genes involved in the response to hypoxia (*Car9*, *Vegfa*, *Pdk1*), amino acid deprivation (*Ass1, Asns)*, and glucose deprivation (*Upp1*) (32–34) (Figure 1L). Consistent with these observations, gene set enrichment analysis (GSEA) indicated enrichment in expression of gene sets related to response to hypoxia and nutrient deprivation in poorly perfused PDAC cells (Figure 1M). Thus, as expected, poorly perfused PDAC cells that exhibit transcriptional hallmarks of metabolic deprivation. In contrast, well perfused PDAC cells exhibited increased expression of genes involved in receptor ligand interaction (*Fgfr1, Epor, Adrb2, Lpar1*) and immune signaling and complement activation (*Cgas, Trex1*) (Figure 1L,N), suggesting there may be significant cell signaling and immunological differences between differently perfused regions of PDAC tumors. Altogether, this analysis indicates that perfusion level substantially changes biology of PDAC cells and identifies marker genes that can be used to differentiate between well and poorly perfused PDAC cells.

With these biomarkers of PDAC perfusion state, we next asked how perfusion is associated with therapeutic response in PDAC patients. To do so, we generated gene sets based on approximately 100 genes whose expression is most significantly associated with poorly perfused PDAC cells. We then identified patients who received chemotherapy as part of their primary PDAC treatment from TCGA (35). These patients were then binned into responders and non-responders. We then performed gene set variation analysis (GSVA) using the poor perfusion gene set and compared relative enrichment between the treatment response groups (Figure 1O). Consistent with prior studies identifying limited perfusion as a biomarker of therapy resistance, tumors enriched in expression of poor perfusion genes had reduced response to therapy and survival time (Figure 1P,Q). Altogether, these studies confirm that PDAC tumors experience significant decreases in perfusion. Furthermore, identification of molecular biomarkers of limited perfusion confirms that the poor perfusion correlates with limited therapy response in patients.

### The TME does not substantially impair chemotherapy exposure in PDAC tumors but strongly mitigates chemotherapy induced cell death

Having established a model that mimics both poor perfusion and therapy resistance observed in human PDAC, we used this model to interrogate how limited perfusion contributes to therapy resistance. One leading model connecting therapy resistance to perfusion is that impaired perfusion decreases drug concentrations in the TME below a therapeutically effective level (20). To assess the extent to which poor perfusion affects intratumoral drug delivery, we treated tumor bearing animals with the chemotherapeutic gemcitabine and collected plasma and tumor interstitial fluid (TIF; the local perfusate of the tumor microenvironment) from animals post-treatment using previously published methods (36). Gemcitabine is rapidly metabolized by deamination and thus displays a relatively transient half-life in the circulation (37). Therefore, we collected plasma and TIF samples at 1 hour post-treatment, approximately when the maximum plasma concentration is reached in mice and humans (12, 37). We then measured the gemcitabine concentration in both the circulation as well as the TIF (Figure 2A) using liquid chromatography-mass spectrometry (Supplemental Figure 2A). Surprisingly, we found that there was not a significant difference in the plasma or TIF concentrations of gemcitabine, with TIF concentrations reaching ∼80 µM (Figure 2B). We also asked if the gemcitabine was able to reach poorly perfused regions within PDAC tumors. We developed an atmospheric pressure-matrix assisted laser desorption/ionization (AP-MALDI) mass spectrometry imaging (MSI) method to measure gemcitabine levels in tumor sections (Supplemental Figure 3A) where perfusion was marked by Hoechst 33342. We observed heterogeneity in gemcitabine distribution but noted both that poorly perfused tumor regions had gemcitabine present (Figure 2C) and that there was no correlation between perfusion and gemcitabine levels (Figure 2D). Thus, there is no substantial impairment of drug delivery into the poorly perfused TME.

**Figure 2:**
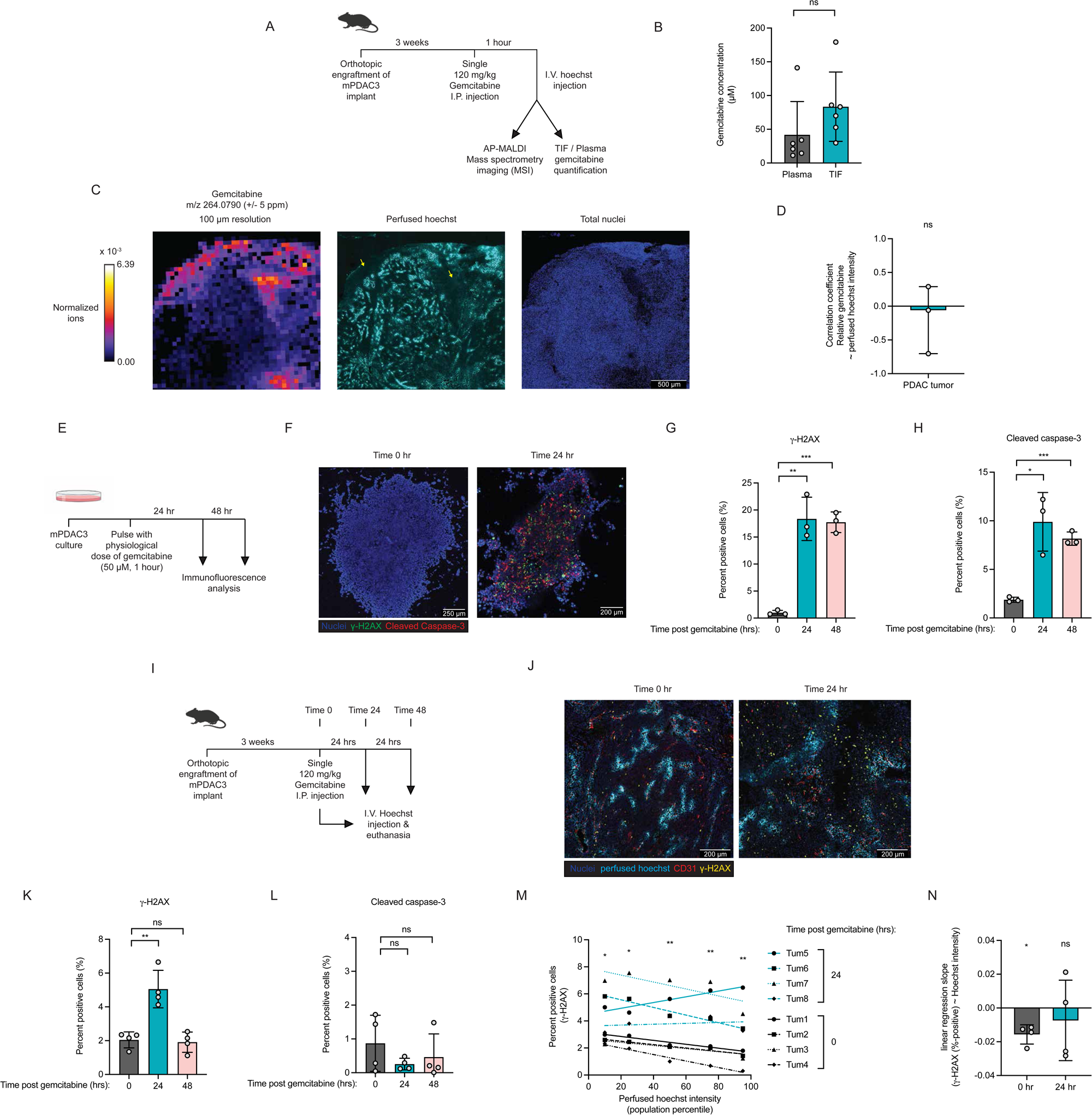
The tumor microenvironment does not substantially impair chemotherapy delivery to PDAC tumors but does disrupt chemotherapy-induced cell death. **(A)** Diagram of workflow measuring gemcitabine concentrations in plasma and tumor interstitial fluid (TIF) samples and measuring gemcitabine spatial distribution in mPDAC3-RPMI tumors by AP-MALDI MSI. **(B)** Levels of gemcitabine in plasma (n = 6) and tumor interstitial fluid (n = 6). **(C)** Representative MSI imaging of gemcitabine and immunofluorescence imaging of perfused Hoechst 33342 (cyan) and total nuclei (blue) for mPDAC3-RPMI tumors in adjacent sections. Yellow arrows indicate poorly perfused regions with high levels of gemcitabine. Scalebar corresponds to 500 µm. **(D)** Correlation coefficient of Hoechst 33342 signal intensity and gemcitabine signal intensity (n = 3). **(E)** Diagram of workflow for treating mPDAC3-RPMI cultures with a physiological pulse of gemcitabine and measuring genotoxic stress (γH2AX labeling) and cell death (cleaved caspase-3 labeling). **(F)** Representative images of γH2AX (green) and cleaved caspase-3 (red) in mPDAC3 *in vitro* cultures following physiological gemcitabine treatment. Scalebar corresponds to 200 µm. **(G)** γH2AX labeling and **(H)** cleaved caspase-3 labeling of mPDAC3-RPMI cultures at 0, 24 and 48 hours post treatment with a physiological dose of gemcitabine (n = 3). **(I)** Diagram of workflow for treatment of mPDAC3-RPMI tumor bearing animals with gemcitabine and Hoechst 33342 prior to immunofluorescence analysis of perfusion (Hoechst 33342 labeling), genotoxic stress (γH2AX labeling) and cell death (cleaved caspase 3 labeling) in tumors. **(J)** Representative immunofluorescence images of mPDAC3 tumors following treatment with gemcitabine. Perfused Hoechst 33342 (cyan), CD31 (red), γH2AX (yellow), total nuclei (blue). Scalebar corresponds to 200 µm. **(K)** γH2AX labeling and **(L)** cleaved caspase-3 labeling of mPDAC3-RPMI tumors at 0, 24 and 48 hours post treatment with gemcitabine (n = 4). **(M)** Percentage of γH2AX labeled tumor cells analyzed by relative intensity of Hoechst 33342 perfusion marker (n = 4). **(N)** Slope of linear regression of the proportion of γH2AX-positive tumor cells as a function of perfused Hoechst 33342 signal intensity from mPDAC3-RPMI tumors at 0 and 24 hours following gemcitabine treatment (n = 4). Statistical significance in B, G, H, K, L and M was determined using 2 sample T-test. For D,N, statistically significant difference from 0 of the correlation coefficients or slopes of regression curves was determined by one sample T-test. * p≤0.05 ** p≤0.01 *** p≤0.001 and **** p≤0.0001.

We next asked if the concentrations of gemcitabine that reach the TME are sufficient for engaging their molecular targets. Gemcitabine is a genotoxic agent, so we assessed if TME levels of gemcitabine can cause DNA damage in two approaches. We treated cultured PDAC cells with a TME-relevant dose of gemcitabine before analyzing these cultures for markers of genotoxic stress (γH2AX) and cell death (cleaved caspase 3) (Figure 2E). Gemcitabine exposure resulted in an increase of DNA damage and cell death (Figure 2F-H). Thus, in cultured cells, exposure to TME physiological levels of gemcitabine causes genotoxic stress and trigger apoptotic death. Second, we measured the ability of gemcitabine treatment to induce DNA damage in PDAC tumors. To do so, we treated tumor bearing mice with a single dose of gemcitabine and recovered tumors for analysis after 24 and 48 hours. Prior to tumor collection, we treated mice with Hoechst 33342 to trace perfusion levels in the tumor (Figure 2I). Across the tumor, we observed a clear induction of DNA damage indicated by γH2AX labeling at 24 hours post gemcitabine treatment, which was fully resolved by 48 hours (Figure 2J,K). We further analyzed how perfusion level impacted the ability of gemcitabine to induce DNA damage. At all perfusion levels, there was a significant increase in the proportion of DNA damaged cells at 24 hours and there was no significant relationship between perfusion level and DNA damage induction (Figure 2M,N). Thus, gemcitabine exerts genotoxic stress even in poorly perfused PDAC tumor regions. Interestingly, we observed no measurable increase in cleaved caspase at any timepoint in gemcitabine treated tumors, despite DNA damage induction (Figure 2L). Altogether, these results indicate that compromised perfusion does not prevent the delivery and action of small molecule drugs like gemcitabine in PDAC. Interestingly, while both PDAC cells in culture and tumors experience DNA damage from physiological levels of gemcitabine, this only results in cell death in cultured PDAC cells, suggesting that the TME may mitigate the efficacy of chemotherapies by suppressing cell death rather than by limiting drug exposure.

### Exposure to TME nutrient levels causes PDAC cells to enter a drug resistant state

We asked how poor perfusion promotes drug resistance if not by limiting drug delivery. Low perfusion has a significant impact on the nutrient conditions present in the TME (27) and recent studies have highlighted regulation of drug response by nutrient availability (27, 38). We therefore hypothesized that TME nutrient availability may contribute to TME-driven drug resistance. To test this hypothesis, we utilized a previously described cell culture media that replicates the physiological concentrations of metabolites within the PDAC TME, <u>T</u>umor Interstitial <u>F</u>luid <u>M</u>edium (TIFM) (28) (Figure 3A). We derived paired mPDAC cell lines (Figure 3B) that are cultured in either standard media (RPMI-1640; mPDAC-RPMI cells) or in TIFM (TIFM; mPDAC-TIFM cells) (28) and treated these cultures with a panel of standard-of-care chemotherapies and experimental targeted therapies and assessed the impact on cell viability (Figure 3C-F). PDAC cells in TIFM were significantly more resistant to all tested therapies (Figure 3C-F and Supplemental Figure 4A-C). TIFM-cultured cells were not only more resistant but maintained proliferative potential and regrew after chemotherapeutic challenge (Supplemental Figure 4D). We also tested if TIFM would promote drug resistance in human PDAC cells. We established patient-derived organoids in TIFM or standard conditions and found TIFM-cultured organoids displayed reduced sensitivity to gemcitabine (Figure 3G). Thus, PDAC cultures established in media with TME levels of nutrients are drug resistant.

**Figure 3:**
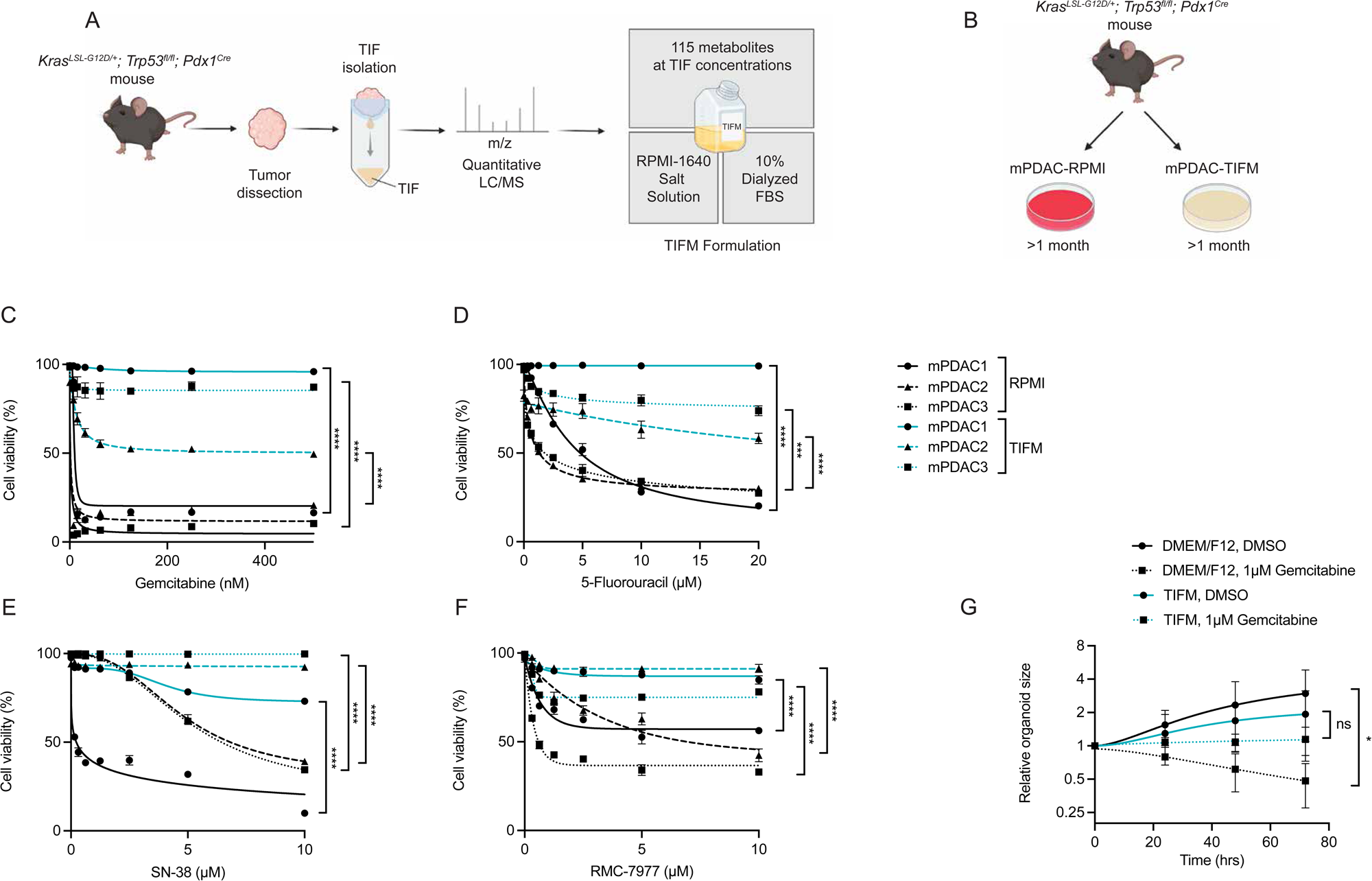
A physiological nutrient model of the tumor microenvironment recapitulates in vivo drug resistance. **(A)** Tumor interstitial fluid composition was determined by quantitative LC/MS and used to generate an RPMI-1640 base media with TME levels of metabolites. **(B)** Diagram of establishing paired murine PDAC cell lines in either RPMI or TIFM medium. **(C-F)** Cell viability of mPDAC-RPMI and mPDAC-TIFM cell lines after 72 hours treatment with the indicated concentration of **(C)** gemcitabine, **(D)** 5-fluorouracil, **(E)** SN-38, or **(F)** RMC-7977. **(G)** Measurement of relative organoid size for TIFM- and standard media-cultured patient-derived organoids over time following treatment with vehicle or gemcitabine (1 µM) (TIFM n = 9, DMEM/F12 n = 6). For C-G, the area under the curve (AUC) was calculated and unpaired t-test was performed to determine significance in the difference of the AUC for the indicated cultures. * p≤0.05 *** p≤0.001 and **** p≤0.0001.

Having established that PDAC cultures derived in TIFM are drug resistant, we next asked if TIFM culture could induce drug resistance in previously drug responsive PDAC cells established in standard laboratory media. We transitioned a panel of murine and human PDAC cell lines and organoids over the course of several weeks to grow in TIFM (RPMI → TIFM culture) (Supplemental Figure 5A) and assessed their response to chemotherapeutic challenge. TIFM-transitioned mPDAC cultures (Supplemental Figure 5B), human PDAC cell lines (Supplemental Figure 5C) and human PDAC organoids (Supplemental Figure 5D) were more resistant to chemotherapeutic exposure than parental cells maintained in standard conditions. Lastly, we asked if resistance arises due to TIFM selecting PDAC subpopulations that are drug resistant. Using lentiviral barcoding (39), we assessed clonal diversity of mPDAC-RPMI cells maintained in RPMI or transitioned to TIFM (Supplemental Figure 5E). Culture of mPDAC cells in RPMI led to a loss of clonal diversity similar to what has been observed in other cancer cell lines (39). Transitioning mPDAC-RPMI cells into TIFM only modestly increased this loss of clonal diversity (Supplemental Figure 5F). Additionally, we observed a significant correlation in clonal abundance in RPMI and TIFM medium conditions (Supplemental Figure 5G). Thus, TIFM does not select for a rare subset of PDAC cells with drug resistance properties but induces this phenotype in previously drug-sensitive PDAC cells. Lastly, we asked if removing PDAC cells from TIFM could reverse the drug resistance phenotype. We cultured mPDAC-TIFM cells in RPMI media (TIFM → RPMI culture) for increasing lengths of time and measured response to chemotherapeutic challenge (Supplemental Figure 5H). At early time points, TIFM → RPMI cultures still maintained a high degree of drug resistance but this was progressively lost (Supplemental Figure 5I). Thus, TIFM-culture conditions lead to drug resistance that is maintained only transiently for a period of days after removal of TME nutrient stress. Collectively, these results suggest that poor perfusion induces a drug resistant state in PDAC cells through chronic exposure of PDAC cells to pathophysiological nutrient levels. However, this state is reversible upon removal of this metabolic challenge.

### TIFM does not cause drug resistance by inhibiting the biochemical activity or cellular response to chemo and targeted therapies

We next sought to understand the mechanisms underlying how TIFM-cultured PDAC cells become resistant to therapeutic challenge. Chemotherapies are typically thought to kill cancer cells in an apoptosis-dependent manner, but at times other cell death mechanisms may be dominant (40). To determine if mPDAC-RPMI cells were killed by apoptosis with chemotherapy exposure, we generated stable knockout cell lines of *Bax* and *Bak1*, which were resistant to chemotherapeutic challenge (Supplemental Figure 6A,B), indicating that indeed chemotherapeutics kills PDAC cells via apoptosis.

PDAC cells in TIFM proliferate slower than cells in standard conditions (28, 41) and have a reduced proportion of cells in S-phase (Supplemental Figure 6C). Many chemotherapeutic agents induce apoptosis only in actively cycling cancer cells (42). Therefore, we first asked if TIFM renders PDAC cells resistant to drug challenge by slowing the proliferative rate of PDAC cells. mPDAC-TIFM cultures remain significantly less sensitive to chemotherapeutic agents even after correcting for differences in cycling rate (43) (Supplemental Figure 6D-H). Lastly, slowing the growth rate of mPDAC-RPMI cultures to that of mPDAC-TIFM cultures through CDK4/6 inhibition did not induce the level of drug resistance observed in mPDAC-TIFM cultures (Supplemental Figure 6I,J). In total, these findings indicate that therapeutic resistance in mPDAC-TIFM cultures cannot be accounted for by slower proliferative kinetics.

Given that slowed proliferation does not explain the ability of TIFM-cultured cells to cope with therapeutic challenge, we next asked which step of the apoptotic cascade (Figure 4A) was impaired in TIFM-cultured PDAC cells. Gemcitabine, 5-Fluorouracil (5-FU) and SN-38 are all drugs that generate DNA damage as part of their anti-cancer effects (44). Therefore, we assessed DNA damage in PDAC cells after drug exposure and found gemcitabine and SN-38 generated similar degrees of DNA damage in TIFM and RPMI cultures, indicating that TIFM culture does not interfere with their activity (Figure 4B,C and Supplemental Figure 7A-C). Gemcitabine induced DNA damage also leads to cell cycle arrest within S-phase and we found that the majority of TIFM-cultured PDAC cells treated with gemcitabine were arrested in S-phase within 24 hours (Figure 4D). In contrast, induction of DNA damage by 5-FU was impaired in TIFM (Supplemental Figure 7D,E). The cytotoxicity of 5-FU is in part mediated by inhibition of thymidylate synthase (TYMS) (45–47) and exogenous thymidine alleviates the cytotoxicity of 5-FU (48) (Supplemental Figure 7F). TIFM contains 2.25 µM thymidine (28), so we asked if exogenous thymidine was causes resistance to 5-FU in TIFM. We found that supplementing mPDAC-RPMI cultures with TIFM levels of thymidine suppressed DNA damage and cell death induced by 5-FU (Supplemental Figure 7E,G). Lastly, we observed that TIFM culture did not impact the ability of KRAS-targeted therapies to inhibit RAS signaling (Supplemental Figure 7H). Together, these results indicate that TIFM culture does not significantly interfere with the biochemical activity of chemo- and targeted therapies in PDAC cells, except for 5-FU. Thus, TIFM-culture promotes drug resistance by enabling cells to persist despite the presence of therapeutic stress, a state which we term therapeutic tolerance.

**Figure 4:**
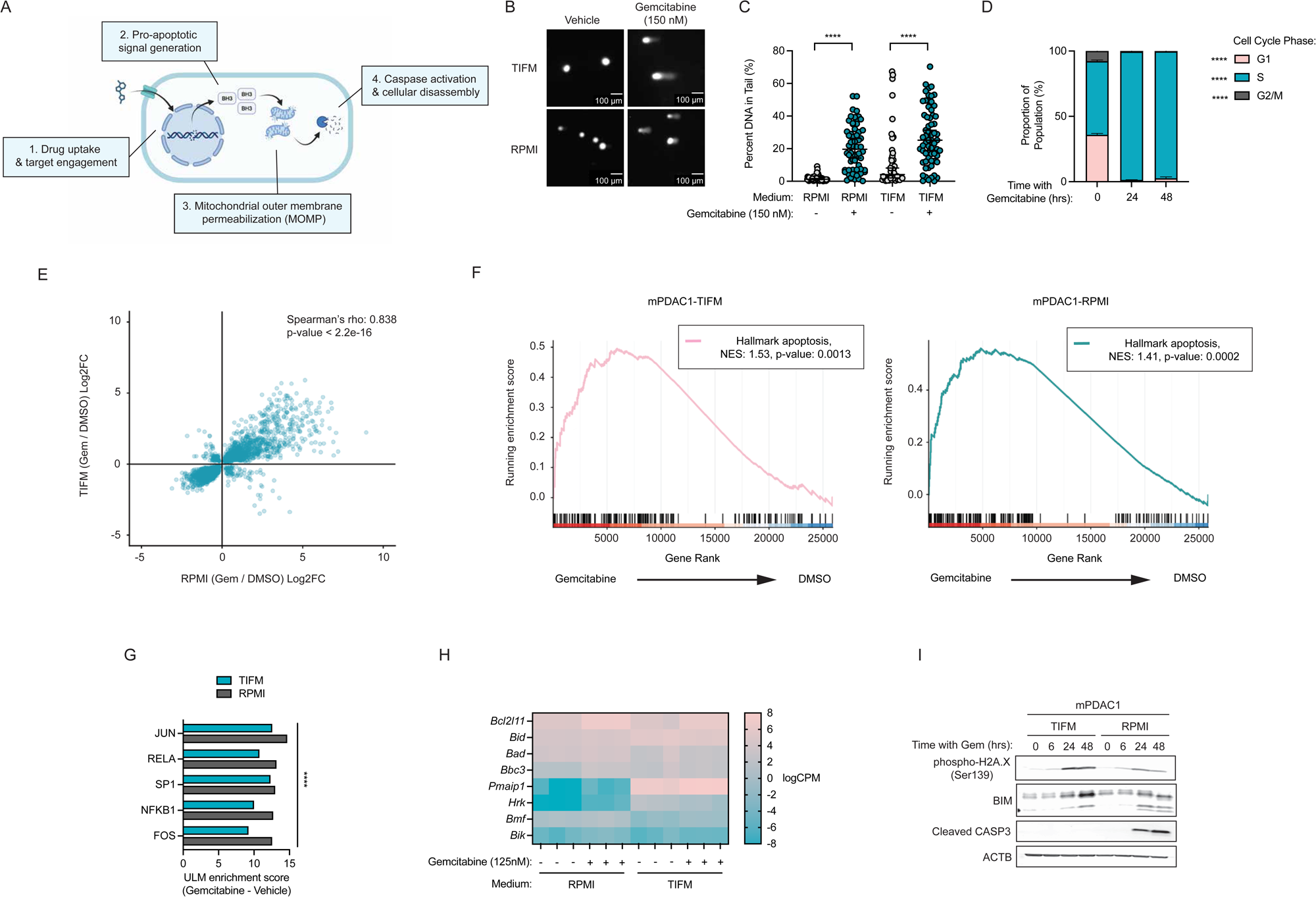
TIFM culture conditions do not interfere with therapeutic target engagement but disrupt execution of apoptosis. **(A)** Diagram of apoptotic cascade following chemotherapy exposure. **(B)** Representative image and **(C)** quantification of alkaline comet assays of mPDAC1-RPMI and mPDAC1-TIFM cultures treated for 18 hours with 150 nM Gemcitabine (RPMI n = 58, TIFM n = 64) or DMSO vehicle (RPMI n = 61, TIFM n = 52). Scalebar corresponds to 100 µm. **(D)** Percentages of mPDAC1-TIFM cells treated with 100 nM gemcitabine for the indicated time periods in G1, S and G2/M phases of the cell cycle (n = 3). **(E)** Correlation of log2 fold change in gene expression with gemcitabine treatment between mPDAC1-RPMI and mPDAC1-TIFM conditions. **(F)** Gene set enrichment analysis for the hallmark apoptosis gene set in mPDAC1-TIFM and mPDAC1-RPMI cultures following treatment with 125nM gemcitabine for 24 hours vs. DMSO control. **(G)** ULM enrichment scores for the top 5 transcription factors with enhanced activity in mPDAC1-RPMI cells treated for 24 hours with 125 nM gemcitabine. The enrichment scores for similarly treated mPDAC1-TIFM cells are displayed in parallel. **(H)** Heatmap of normalized count values for known pro-apoptotic BH3-only genes following RNA-seq analysis of mPDAC1-RPMI and mPDAC1-TIFM cells treated with 125 nM gemcitabine or vehicle for 24 hours. **(I)** Immunoblots for γH2AX, BIM, and cleaved caspase-3 in mPDAC1-RPMI and mPDAC1-TIFM cells treated with 150 nM gemcitabine over time at the indicated timepoints. Statistically significant differences in C,D were determined by two-sample T-test. For E, the correlation coefficient was determined using spearman’s rank correlation coefficient method. For G, all transcription factors displayed had significant enrichment with a p-value of less than p=0.0001 after multiple tests correction. **** p≤0.0001.

Given that TIFM-culture does not prevent the biochemical activity of PDAC therapeutics, we next asked if TIFM-culture alters downstream stress-induced apoptotic signaling. Gemcitabine treatment induces a similar transcriptional response in PDAC cells in either RPMI or TIFM (Figure 4E), including a similar increase in JUN and FOS transcriptional programs, two transcription factors with critical roles in promoting cell death signaling (49, 50), in response to gemcitabine treatment (Figure 4G). This resulted in similar upregulation of apoptotic gene expression (Figure 4F) between culture conditions. We also observed similar regulation of pro-apoptotic BCL2 family genes in PDAC cells cultured in both conditions, with gemcitabine inducing strong upregulation of *Bcl2l11*, encoding the apoptotic activator protein BIM (Figure 4H). We also observed significant upregulation of BIM by immunoblotting in both TIFM- and RPMI-cultured PDAC cells upon chemotherapy exposure (Figure 4I). Collectively, these results indicate that metabolic stress in the poorly perfused TME does not substantially impair the induction of pro-apoptotic signals in response to therapy challenge.

### TIFM causes drug resistance by disrupting execution of apoptosis in a BCL-XL-dependent manner

The last step in therapy-induced apoptosis is mitochondrial outer membrane permeabilization (MOMP) and activation of executioner caspases (Figure 4A). We observed caspase activity in chemotherapy treated PDAC cells cultured in RPMI but not in TIFM (Figure 4I), suggesting a defect in execution of apoptosis upon therapeutic challenge. We next asked if this defect resulted from a failure in MOMP. We assessed MOMP induction using three approaches. First, using a genetically-encoded sensor of mitochondrial integrity (51) we found that MOMP readily occurred in mPDAC-RPMI cultures challenged with gemcitabine, but not mPDAC-TIFM cultures (Figure 5A,B). Second, MOMP activation is triggered by the accumulation and oligomerization of pore-forming proteins such as BAK and BAX at the mitochondria but which are otherwise continuously retrotranslocated back to the cytosol under homeostatic conditions (52). We observed that BAX translocated from the cytosol to the mitochondria with gemcitabine treatment in RPMI-cultured by not TIFM-cultured PDAC cells (Figure 5C). Lastly, we assessed MOMP propensity in PDAC cells using BH3 profiling (53). Relative to PDAC cells cultured in RPMI, we observed limited mitochondrial depolarization in response to BIM peptide in TIFM-cultured PDAC cells, indicative of reduced MOMP propensity under these conditions (Figure 5D).

**Figure 5:**
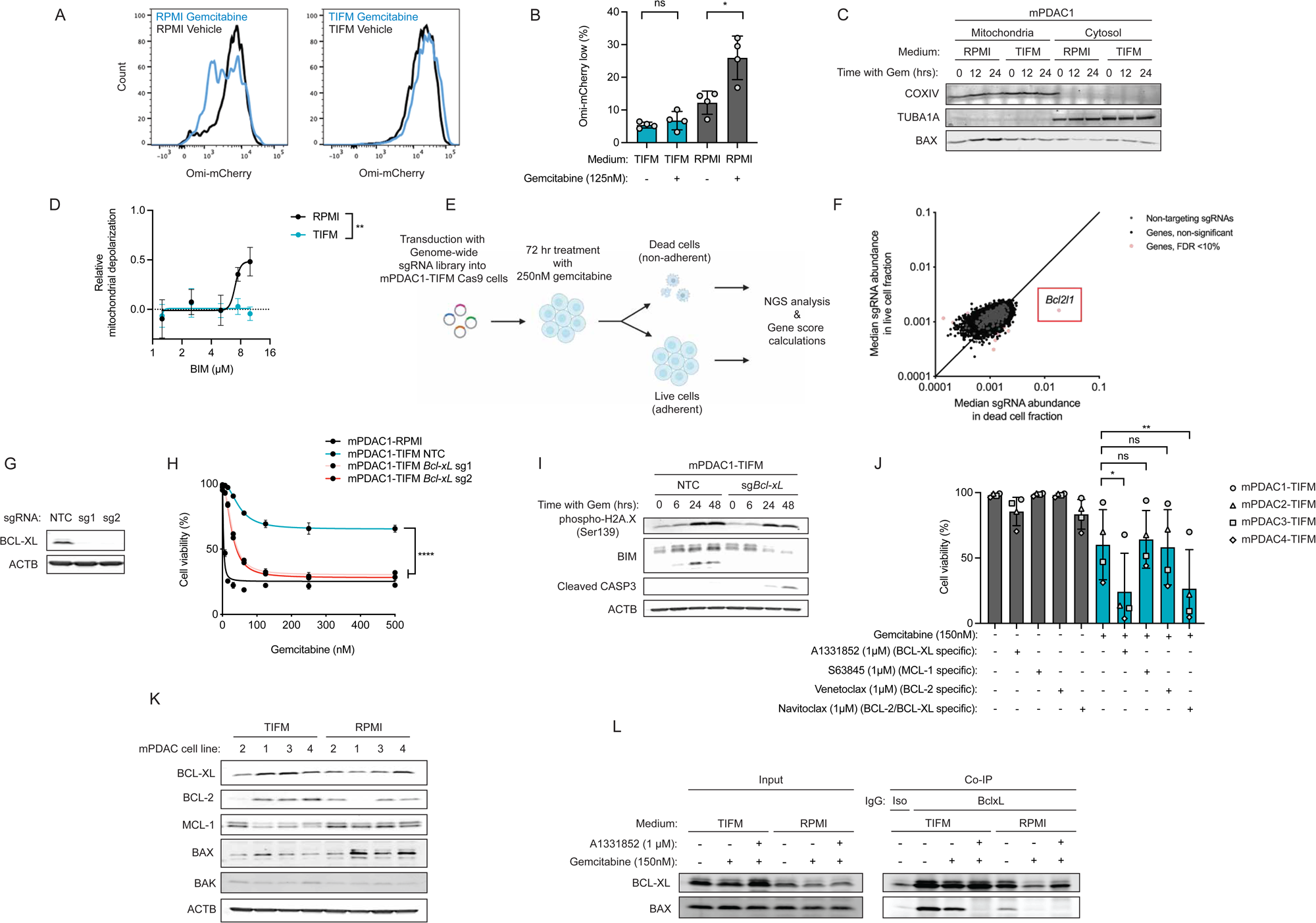
TIFM culture conditions suppress apoptotic priming via increased BCL-XL activity. **(A)** Fluorescence intensity of the Omi-mCherry reporter in mPDAC1-RPMI and mPDAC1-TIFM cells treated with either gemcitabine or vehicle. **(B)** Percent of Omi-mCherry low cells (post-MOMP) in mPDAC cells cultured with indicated medium and drug (n = 4). **(C)** Immunoblots of BAX in mitochondrial and cytosolic isolates from mPDAC1-RPMI and mPDAC1-TIFM cells treated with 250 nM gemcitabine at the indicated timepoints. **(D)** Relative mitochondrial depolarization of mPDAC1-RPMI and mPDAC1-TIFM cells in response to treatment with BIM peptide at the indicated concentrations (n = 3). **(E)** Diagram of Death-Seq CRISPR screen to detect genes sensitizing mPDAC-TIFM cultures to gemcitabine. **(F)** Median sgRNA abundance in live and dead cell fractions following gemcitabine treatment. Genes with statistically significant enrichments in live or dead cell fractions based on MAGeCK MLE analysis are highlighted in pink. *Bcl2l1* enrichment in dead cell fraction highlighted. **(G)** Immunoblot confirmation of BCL-XL knockout in mPDAC1-TIFM cells. **(H)** Cell viability of BCL-XL knockout or control mPDAC1-TIFM cells following treatment with the indicated concentrations of gemcitabine for 72 hours (n = 3). **(I)** Immunoblots for γH2AX, BIM, and cleaved caspase-3 in BCL-XL knockout or NTC mPDAC1-TIFM cells following treatment with 150 nM Gemcitabine at the indicated timepoints. **(J)** Cell viability of mPDAC1-TIFM cells following treatment with the indicated compounds for 72 hours (Gemcitabine, 150 nM), (A1331852 1 µM), (S63845 1 µM), (Venetoclax 1 µM), (Navitoclax 1 µM). Individual datapoints reflect the mean viability of 3 replicates for a single cell line (mPDAC1-4), as indicated in the figure legend (n = 4 cell lines). **(K)** Immunoblots for apoptotic regulators in mPDAC-RPMI and mPDAC-TIFM cell lines. **(M)** Immunoblots for BCL-XL and BAX following immunoprecipitation for BCL-XL in mPDAC1-TIFM or mPDAC1-RPMI cell lysates following treatment with gemcitabine (125 nM) and A1331852 (1 µM) as indicated for 36 hours. Statistically significant differences for B, J, L were determined by two-sample T-test. For D,H, the area under the curve (AUC) was calculated and unpaired t-test was performed to determine significance in the difference of AUCs for the indicated cultures. For F, adjusted p-values were calculated using Benjamini-Hotchberg (BH) false discovery rate correction and genes highlighted in pink. * p≤0.05 ** p≤0.01 *** p≤0.001 and **** p≤0.0001.

Altogether, these data indicate that TME metabolic conditions limit MOMP and execution of apoptosis downstream of therapy induced stress and pro-death signaling. We next sought to understand how MOMP is suppressed in TIFM-cultured PDAC cells. Pooled genetic screens that positively select for dead cells have proven especially suited in identifying suppressors of cell death (54, 55). Therefore, we performed a Death-seq screen (54), where we mutagenized mPDAC1-TIFM cultures with a genome wide CRISPR-library and assessed enrichment of guides in a purified dead cell population compared to remaining viable cells after treatment with gemcitabine (Figure 5E). Guide RNAs targeting *Bcl2l1*, the gene encoding for the anti-apoptotic protein BCL-XL, were by far the most enriched guides in the dead cell population (Figure 5F). CRISPR knockout *Bcl-xL* from mPDAC cells confirmed a critical role for BCL-XL in TIFM-driven chemoresistance and apoptosis suppression (Figure 5G-I). Additionally, pharmacological perturbation of BCL-XL, but not other anti-apoptotic proteins, using BH3-mimetic compounds consistently enhanced the ability of chemotherapy to induce cell death in TIFM-cultured PDAC cells (Figure 5J). Thus, BCL-XL is critical to TIFM-induced drug resistance.

Lastly, we asked how TIFM-induces BCL-XL-dependent drug resistance. We blotted for BCL-XL in mPDAC-TIFM and mPDAC-RPMI cultures and did not observe consistent upregulation of BCL-XL in drug resistant TIFM-cultures (Figure 5K). BCL-XL normally functions by binding to pro-apoptotic effector proteins such as BAX and BAK, preventing their activation under homeostatic conditions (56). We found that the BCL-XL interaction with BAX was not significantly impacted by gemcitabine treatment TIFM-cultured PDAC cells but was completely lost upon gemcitabine treatment in RPMI-cultured PDAC cells (Figure 5L). The interaction was fully lost in both culture conditions upon treatment with BCL-XL-targeting BH3 mimetics (Figure 5L). This finding is consistent with reduced apoptotic priming in TIFM-cultured cells being driven by enhanced BCL-XL function rather than increased BCL-XL levels. Altogether, this genome-wide search identified BCL-XL as a critical anti-apoptotic protein which PDAC cells require for therapeutic tolerance.

### Chronic nutrient stress reduces apoptotic priming and therapy response in PDAC cells

We next asked which metabolic differences between TIFM and standard media decrease apoptotic potential and therapy response in PDAC cells. We focused on amino acid deprivation as genes involved in response to amino acid deprivation are enriched in poorly perfused PDAC cells (Figure 1M) and in TIFM (28). Furthermore, expression of these genes correlates with poor outcomes in PDAC patients (Figure 6A). Of the amino acids, we focused on arginine starvation as we previously identified myeloid arginase-1-driven arginine starvation as a major metabolic stress in the PDAC TME (28, 36). Furthermore, arginase-1 expression is particularly prevalent in poorly perfused regions of PDAC (Figure 6B-D) suggesting this metabolic stress is relevant in poorly perfused PDAC regions. Lastly, arginine levels are substantially lower in TIFM compared to standard media (28), and we found supplementation of arginine to TIFM resolves the amino acid deprivation transcriptional signature in PDAC cells (Figure 6E). Altogether, this results suggested that arginine starvation could be a critical driver of perfusion stresses in PDAC tumors and TIFM.

**Figure 6:**
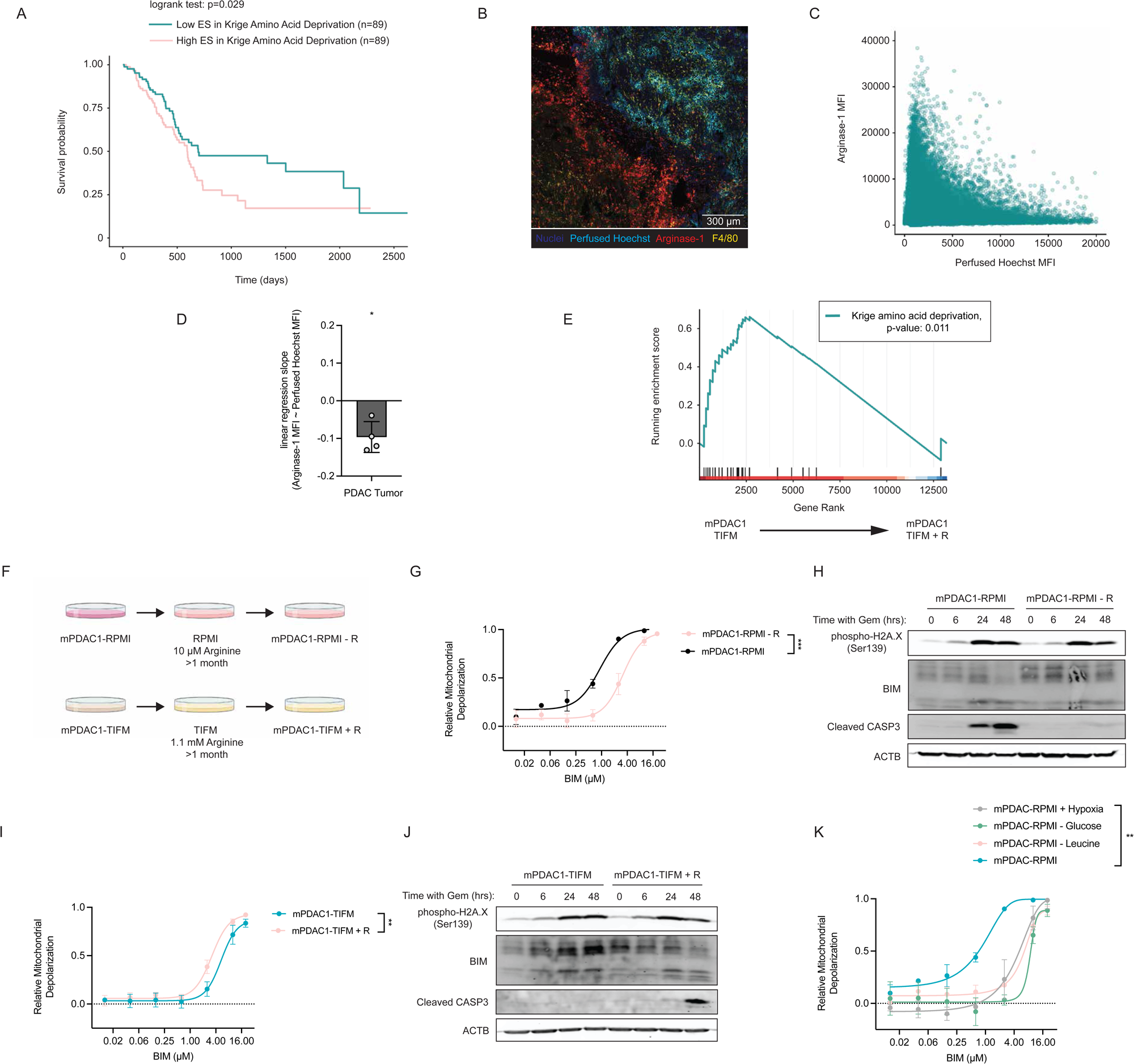
Chronic amino acid stress reduces apoptotic priming and therapy response in PDAC. **(A)** TGCA PDAC patients were stratified into cohorts with high (n = 89) or low (n =89) GSVA enrichment for the Krige Amino Acid Deprivation gene set and Kaplan Meier survival analysis was performed. **(B)** Representative image of Hoechst 33342 staining (cyan) and arginase-1 levels (red), F4/80 (yellow), and total nuclei (blue) in a PDAC tumor. **(C)** Scatterplot of Hoechst 33342 staining and arginase-1 levels in PDAC tumors (n = 30,000 cells). **(D)** Slope of linear regression of arginase-1 levels as a function of Hoechst 33342 labeling in PDAC tumors (n = 4). **(E)** Gene set enrichment analysis of the Krige Amino Acid Deprivation gene set for mPDAC1-TIFM vs. mPDAC1-TIFM + R cultures. **(F)** Diagram of establishment of arginine-starved mPDAC-RPMI cultures (mPDAC1-RPMI – R) and arginine supplemented mPDAC1-TIFM cultures (mPDAC1-TIFM + R). **(G)** Relative mitochondrial depolarization of mPDAC1-RPMI and mPDAC1-RPMI - R cells with BIM peptide treatment at the indicated concentrations (n = 3). **(H)** Immunoblots for γH2AX, BIM, and cleaved caspase-3 in mPDAC1-RPMI and mPDAC1-RPMI - R cells following treatment with 150 nM gemcitabine at the indicated timepoints. **(I)** Relative mitochondrial depolarization of mPDAC1-TIFM and mPDAC1-TIFM + R cells with treatment of BIM peptide at the indicated concentrations (n = 3). **(J)** Immunoblots for γH2AX, BIM, and cleaved caspase-3 in mPDAC1-TIFM and mPDAC1-TIFM + R cells following treatment with 150 nM gemcitabine at the indicated timepoints. **(K)** Relative mitochondrial depolarization following treatment with BIM peptide of mPDAC1-RPMI and cultures adapted to the indicated nutrient stress for 4-weeks (n = 3). Statistical significance was determined in A by log-rank test. For D, statistically significant difference from 0 for the slopes of regression curves was determined by one sample T-test. For G,I,K the area under the curve (AUC) was calculated and unpaired t-test was performed to determine significance in the difference of AUCs for the indicated cultures. * p≤0.05 ** p≤0.01 and *** p≤0.001.

Therefore, we asked if arginine restriction in TIFM caused drug resistance in PDAC cells. We evaluated apoptotic priming and caspase activation following chemotherapy treatment in RPMI-cultured PDAC cells that were adapted to grow in levels of arginine present in the PDAC TME (10 µM) (28, 36) for over a month (Figure 6F). Arginine restriction resulted in suppressed apoptotic priming (Figure 6G) and drug-induced caspase activation (Figure 6H) in PDAC cells. Conversely, arginine supplemented PDAC TIFM cultures (Figure 6F) increased apoptotic priming (Figure 6I) and caspase activity upon chemotherapy exposure (Figure 6J). Thus, these results indicate that chronic TME arginine restriction can reduce apoptotic priming and PDAC cell killing by chemotherapies.

Poor perfusion impacts the availability of multiple nutrient levels within the TME beyond arginine. Therefore, we additionally asked whether restriction of other nutrients would affect apoptotic priming in PDAC. We established cultured drug-sensitive RPMI-cultured PDAC cells in media with the following metabolic stresses: low leucine (10 µM), low glucose (1 mM) or hypoxia (0.5% oxygen). We found that all these conditions resulted in suppression of apoptotic potential relative to standard culture conditions (Figure 6K). Thus, prolonged exposure to multiple other metabolic stresses in the PDAC TME induces the same apoptosis resistant state in PDAC cells, suggesting that poor perfusion could induce apoptosis resistance via multiple metabolic stresses.

### The TME dampens apoptotic priming in PDAC

These results suggest that prolonged contact with the stressful metabolic conditions within the poorly perfused TME give rise to PDAC cells with dampened apoptotic priming that are poised to survive chemotherapeutic challenge. To test this hypothesis, while no methods for assessing apoptotic priming *in situ* exist, given that TME nutrient stress induces dampened apoptotic priming and drug resistance that persists for a period of days after removal of this stress, we determined if PDAC cells recently isolated from the TME would have reduced apoptotic priming and increased drug resistance. We established orthotopic allograft PDAC tumors or maintained the same cells in standard culture conditions. We isolated PDAC cells from orthotopic tumors and measured their apoptotic priming and response to chemotherapeutic challenge after both a short period (5 days) or prolonged period in standard culture (4 weeks) (Figure 7A). These results were compared to paired cultures that were maintained in RPMI (Figure 7A). In PDAC cells removed from the TME for 5 days, we observed significant suppression of apoptotic priming (Figure 7B) and response to chemotherapeutic agents compared to the paired culture maintained in RPMI (Figure 7C,D). In contrast, PDAC cells removed the TME for 4 weeks no longer had a significant change in apoptotic priming (Figure 7E) or response to therapeutic challenge (Figure 7F,G). Thus, as with TIFM-exposure, TME contact suppresses the apoptotic priming of PDAC cells, decreasing their sensitivity to chemotherapeutic agents. However, as with TIFM-exposure, TME-exposure does not result in permanent reprogramming of PDAC cells to a drug-resistant state and this state is lost upon prolonged culture of PDAC cells in standard laboratory conditions. Altogether, these results suggest that PDAC cells in the TME are in a state of dampened apoptotic priming that enables them to tolerate therapeutic challenge.

**Figure 7:**
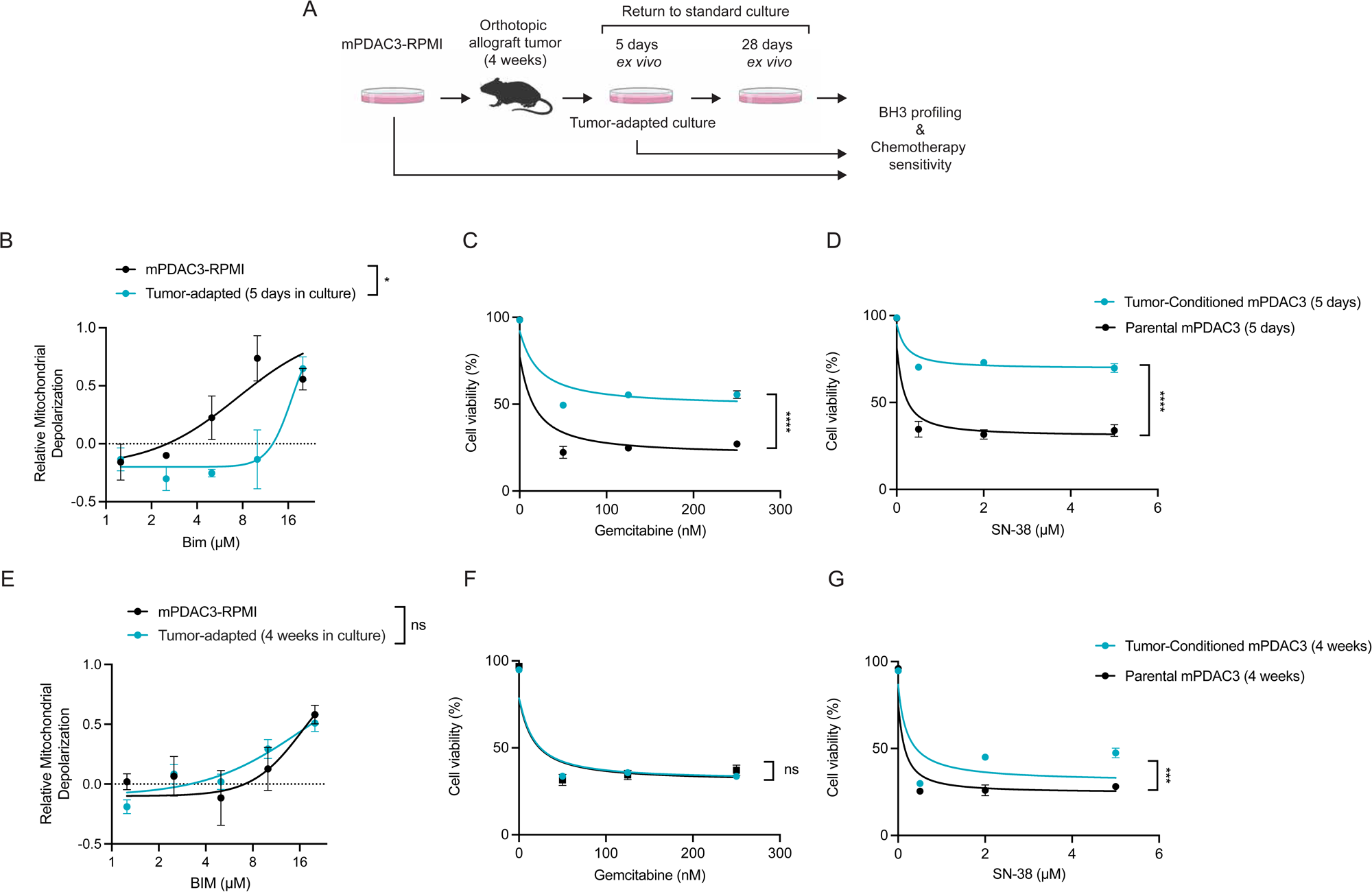
The TME suppresses apoptotic priming and chemotherapy sensitivity in PDAC. **(A)** Diagram of generating mPDAC3-RPMI tumor-adapted cells and paired cultures maintained in standard conditions. **(B)** Relative mitochondrial depolarization of mPDAC1-RPMI and tumor-adapted cultures following 5 days of ex vivo culture in response to treatment with BIM peptide at the indicated concentrations (n = 3). Cell viability of mPDAC1-RPMI and tumor-adapted cultures following 5 days of ex vivo culture with treatment for 72 hours with **(C)** gemcitabine and **(D)** SN-38 measured by propidium iodide exclusion (n = 3). **(E)** Relative mitochondrial depolarization of mPDAC1-RPMI and tumor-adapted cultures following 4 weeks of ex vivo culture in response to treatment with BIM peptide at the indicated concentrations (n = 3). Cell viability of mPDAC1-RPMI and tumor-adapted cultures following 4 weeks of ex vivo culture after treatment for 72 hours with **(F)** gemcitabine and **(G)** SN-38 measured by propidium iodide exclusion (n = 3). For B-G, the area under the curve (AUC) was calculated and unpaired t-test was performed to determine significance in the difference of AUCs for the indicated cultures. * p≤0.05 *** p≤0.001 and **** p≤0.0001.

## Discussion

### Drug tolerance rather than impaired drug delivery drives therapy resistance in poorly perfused PDAC tumors

Poor perfusion and drug resistance are hallmark of PDAC. Poor perfusion correlates with limited therapeutic resistance (18, 19), suggesting a link between these features of PDAC. Thus, understanding how limited perfusion contributes to therapy resistance will be critical to improving patient outcomes. One proposed mechanism by which limited perfusion causes drug resistance is by impairing the delivery of therapeutic agents to the tumor (20). We used interstitial fluid analysis and imaging studies to measure the amount and distribution of a therapeutic agent and its biochemical activity in a mouse model of poorly perfused and therapy resistance PDAC. We found that the widely used chemotherapy gemcitabine does accumulate at therapeutic levels across the PDAC TME and triggers genotoxic stress even in the most poorly perfused tumor regions. This suggested that poor perfusion does not cause drug resistance by limiting drug delivery. In line with these findings, PDAC cells removed from tumors and drug treated ex vivo initially have substantially reduced sensitivity to chemotherapeutic agents before reverting to a drug sensitive state. Altogether, these findings suggest that the poorly perfused TME reduces drug efficacy by imprinting a drug resistant state on PDAC cells, rather than by reducing drug exposure. These findings have important implications for ongoing efforts to improve PDAC therapeutic responses by increasing intratumor drug delivery (21–23, 57) and suggest that these approaches may not target the most critical barrier to therapy efficacy in this disease.

We next sought to understand how the poorly perfusion imprints the drug resistant state in PDAC. We found that culture in TIFM, which recapitulates the nutrient levels in the TME, reprograms PDAC cells to be resistant to therapeutic treatment by dampening apoptotic priming such that PDAC cells tolerate therapeutic challenge. We additionally observed evidence of drug tolerance in animal tumors, where chemotherapy treated tumors exhibited evidence of on-target activity but not commitment to cell death. Furthermore, PDAC cells directly isolated from tumors were therapy tolerant and exhibited reduced apoptotic priming. Altogether, this suggests that pathophysiological nutrient levels in the PDAC TME drive therapy resistance in PDAC by imprinting a therapy tolerant state. Importantly, this mode of drug resistance appears operative in human PDAC. We found that human PDAC cells and organoids are reprogrammed to a drug resistant state by TIFM culture and analysis of chemo- and radiotherapy treated patient PDAC tumors indicates the presence of therapy tolerant persister cells (58, 59). Mechanistically, therapeutic tolerance is dependent upon BCL-XL. Importantly, we do not understand the molecular basis for how BCL-XL is regulated by the TME or TIFM to induce therapy tolerance. Upregulation of BCL-XL was not consistently observed in our studies of drug tolerant PDAC cells, arguing that expression levels of this anti-apoptotic protein alone are not fundamental to the acquisition of this cell state. Instead, we hypothesize there must be another mode of regulation of BCL-XL regulation that promotes the limited apoptotic potential and drug tolerance of these cells, which future studies should address. Nevertheless, our findings that BCL-XL-mediated apoptotic resistance is key to drug tolerance in PDAC is consistent with recent findings that targeting BCL-XL can improve therapeutic outcomes in animal models PDAC (60–64) and suggest that targeting critical regulators of drug tolerance could improve efficacy of PDAC treatments.

While our studies largely focused on PDAC response to chemotherapies, the mechanism of drug tolerance by dampening apoptotic priming is likely to mitigate the response to any therapy that relies on engaging this mode of cell death. Notably, RAS targeted therapies exert their efficacy in part via apoptosis (6, 7, 65). We found that TIFM-cultured PDAC cells exhibited tolerance to RAS targeted therapies. Furthermore, drug tolerant persister cells lead to relapse of murine PDAC tumors treated with RAS targeted therapies (66). Thus, future studies determining if TME suppression of apoptosis contributes to targeted therapy resistance are warranted.

### The drug tolerant PDAC cell state is imprinted by the TME but not maintained in standard culture conditions

An interesting feature of TME-imprinted drug tolerance is the transient nature of this phenotype. PDAC cells isolated from tumors or cultured in TIFM exhibited drug tolerance only for a period of days after isolation from the tumor or removal from TIFM. This suggests a temporary cellular “memory” of the TME. Future studies understanding the molecular basis of this memory will be necessary to understand how drug tolerance is imprinted by nutrient levels in the TME. Understanding this imprinting could lead to new approaches to limit the development of drug tolerance to improve the efficacy of therapies in PDAC treatment.

The finding that standard culture models cannot maintain PDAC cells in a physiologically relevant drug tolerant state also has important implications for the use of these models in preclinical drug efficacy studies. Culture models have proven useful for identifying genetic determinants of therapy response (67–70), which has been key in the development of precision medicine. However, tumor response to cytotoxic and targeted therapies are often dampened compared to what is predicted from culture models (71). Many explanations for these discrepancies have been put forward including differences in growth rate, phenotypic heterogeneity and differences in drug exposure between cancer cells in culture and in tumors (71). Our data suggests an important additional driver of this disconnect between culture models and tumors, namely the increased apoptotic priming of cancer cells in standard culture conditions. Apoptotic priming is a critical determinant of therapy response (72), so the increase of apoptotic priming that occurs in cancer cells in standard culture conditions could explain why therapies that induce apoptosis often have stronger effects in culture models than in tumors. This standard culture media-induced shift in apoptotic priming could limit the predictive power of culture models when used both for drug discovery and as ex vivo precision medicine platforms (73–75). Thus, development of culture systems, such as TIFM, that better represent the tumor microenvironment and the changes in cell state that the TME confers will be critical to improve the predictive power of ex vivo cancer models.

### TME metabolic stress induces a drug tolerant PDAC cell state

PDAC is poorly perfused due to hypovascularity (14) and impaired vascular functionality (12, 16). These features of PDAC tumors result in abnormal nutrient levels in the TME (27). Nutrient limitation and metabolic stress select cells with diminished apoptotic potential (76–78) and are linked to drug tolerance in cancer (79). This led us to hypothesize that nutrient stress in the TME could push cells into a state of decreased apoptotic potential resulting in cross-tolerance to apoptosis-inducing therapies. Our finding that TIFM culture induces a drug tolerant state indeed demonstrates that TME metabolic stress is an important contributor to limited therapy response in PDAC. Further, we were able to identify amino acid deprivation, a well-known metabolic stress in the PDAC TME (27), as a driver of this phenotype. Interestingly, analysis identified a gene signature associated with amino acid stresses that correlates with poor patient outcomes (80), consistent with our findings that amino acid deprivation reprograms PDAC to a therapy resistant state. Thus, amino acid stress appears to be an important TME factor pushing PDAC cells to a drug tolerant state. Importantly, these studies do not rule out the contribution of other metabolic stresses in promoting drug resistance in PDAC. Indeed, we found that other TME-relevant metabolic stresses such as hypoxia and glucose deprivation can push PDAC cells to state of dampened apoptotic priming, which is consistent with studies showing that chronic exposure to these environments can trigger drug resistance in PDAC cells (78, 81, 82). Thus, we speculate that multiple metabolic stresses that arise in poorly perfused PDACs all converge on pushing PDAC cells to a drug resistant state. Future studies identifying how TME metabolic stresses rewire PDAC cells to trigger drug tolerance will provide important insight into therapy resistance in PDAC.

## Material and methods

Further information can be found in Supplemental methods.

### Cell lines and culture

Murine PDAC lines were derived from *Kras^LSL-G12D/+^; Trp53^fl/fl^;Rosa26 ^tm1(EYFP)Cos^; Pdx1^Cre^* (mPDAC1-3) and *Kras^FSF-G12D/+^;Trp53^frt/frt^;Pdx1^Flp^* (mPDAC4) autochthonous mouse models of PDAC as previously described (28, 41). Human cancer cell lines, ASPC1 and 8988T, were generously donated from the lab of Dr. Matthew Vander Heiden. Use of cancer cell lines was approved by the Institutional Biosafety Committee (IBC 1560). All cancer cell lines were tested quarterly for mycoplasma contamination using the Mycoalert Mycoplasma Detection Kit (Lonza, LT07-318). All cells were cultured in Heracell Vios 160i incubators (Thermo Fisher) at 37°C and 5% CO_2_. Cell lines were routinely maintained in RPMI-1640 or TIFM supplemented with 10% dialyzed FBS (Gibco, 26400-044).

### Animal experiments

Animal experiments were approved by the University of Chicago Institutional Animal Care and Use Committee (IACUC, Protocol #72587) and performed in strict accordance with the Guide for the Care and Use of Laboratory Animals of the National Institutes of Health. Mice were housed in a pathogen-free animal facility at the University of Chicago with a 12 hour light / 12 hour dark cycle. Humidity is maintained between 30–70% humidity and temperature is maintained at 68–74 °F in the facility.

### Generation of orthotopic allograft PDAC tumors

Female C57BL6J mice 8–12 weeks of age were obtained from Jackson Laboratories (Strain #000664). mPDAC3-RPMI cells were resuspended in sterile serum-free RPMI and Reduced Growth Factor Basement Membrane Extract (R&D Biosystems, 3433-010-01) at a final concentration of 5.6 mg/ml. To generate orthotopic tumors, 100,000 cells were injected as a 20 µL Reduced Growth Factor Basement Membrane Extract:PDAC cell mixture into the splenic lobe of the pancreas of anesthetized mice as previously described (83). Following implantation, mice were regularly monitored for changes in behavior, body weight, and tumor presence by abdominal palpation.

### Human samples regulation

Human PDAC samples were obtained under approval by the Institutional Review Board at the University of Chicago (20-1683).

### Derivation of patient-derived organoids in TIFM medium

Human pancreatic ductal adenocarcinoma (PDAC) biopsy specimens were obtained under institutional review board–approved protocols with informed patient consent. Fresh tissue was transferred into digestion medium consisting of Advanced DMEM/F12 (Gibco, 12634010) supplemented with 1× penicillin/streptomycin (Gibco, 15140163), 1× GlutaMAX (Gibco, 35050061), 10 mM HEPES (Gibco, 15630080), and 100 µg/mL Primocin (InvivoGen, ant-pm-05), containing 0.125 mg/mL collagenase IV (Sigma-Aldrich, C9407) and 0.125 mg/mL Dispase II (Sigma-Aldrich, D4693). Tissue was minced until the pieces were barely visible by eye, then transferred to 15 mL conical tubes containing digestion medium and incubated for exactly 30 minutes at 37 °C on an orbital shaker set to 100 rpm. Following digestion, the suspension was passed through a pre-wetted 100 µm cell strainer (Falcon, 352360) using a pipette to facilitate passage, and the filtrate was collected into a new tube. The suspension was centrifuged at 500 × g for 5 min at 4 °C, and the pellet was washed in Advanced DMEM/F12 with 1% FBS (Corning, 35-015-CV).

Cells were embedded in pure Cultrex UltiMatrix Reduced Growth Factor Basement Membrane Extract (Bio-Techne, BME001) and plated in two 35 µL domes per well in pre-warmed 24-well plates (Greiner, 662102). Plates were inverted and incubated at 37 °C for 20–30 min before overlaying with 500 µL of complete PDAC organoid medium. The medium comprised TIFM medium supplemented with 10.5 µM Y-27632 (Cayman Chemical, 10005583), 300–360 ng/mL recombinant human Noggin (ImmunoPrecise Antibodies, N002), 800–880 ng/mL recombinant human R-spondin (ImmunoPrecise Antibodies, R001), 1× B27 supplement (Gibco, 17504044), 5 mM nicotinamide (Thermo Scientific, 128271000), 100 µg/mL Primocin (InvivoGen, ant-pm-05), 50 ng/mL recombinant human EGF (PeproTech, AF-100-15), 10 ng/mL heat-stable recombinant human FGF-10 (Gibco, PHG0371), 500 nM A83-01 (BioGems, 9094360), and 100 nM gastrin (Sigma-Aldrich, G9145).

Cultures were maintained at 37 °C in a humidified incubator with 5% CO_2_, with medium replaced every 2–3 days. Organoids were passaged when the average diameter reached 200–300 µm, using TrypLE with Y-27632, and re-embedded in fresh UltiMatrix domes as above.

### Patient-derived organoid derivation in standard organoid conditions

Human PDAC biopsy specimens were obtained from surgical resections under institutional review board–approved protocols with informed patient consent. Organoids were isolated as previously described (84). Briefly, surgical tumor pieces were minced into 1 mm fragments with a sterile scalpel and enzymatically digested with Accutase (Sigma-Aldrich, A6964) for 30 min at 37 °C on an orbital shaker. Cells were then embedded into 50 µL domes of Cultrex Reduced Growth Factor Basement Membrane Extract, Type 2, Pathclear (R&D Systems, 3533-001-02) into 24-well plates and covered with human complete medium (hCPLT) containing Advanced DMEM/F-12 (Gibco, 12634028), HEPES (Gibco, 15630-080), GlutaMAX (Gibco, 35050061), Primocin (Invivogen, ant-pm-2), WNT3A conditioned media, R-spondin1 conditioned media, B27 supplement (Gibco, 17504-001), Nicotinamide (Sigma-Aldrich, N0636), N-acetylcysteine (Sigma-Aldrich, A9165), mouse Noggin (Gibco, AF-250-38), human EGF (Gibco, AF100-15), human FGF (Gibco, AF-100-26), human Gastrin I (Tocris Bioscience, 3006), A 83-01 (Tocris Bioscience, 2939) and Y-27632 Rho Kinase inhibitor (Sigma-Aldrich, Y0503). Cultures were maintained at 37 °C in a humidified incubator with 5% CO_2_. The culture medium was replaced regularly, and organoids were passaged for cell expansion approximately every 5 to 14 days depending on cell density and growth rate.

### Statistical analysis

Graphpad Prism was used for all statistical comparisons for all numerical comparisons between groups as well as computing areas under curves for drug treatment and BH3 profiling experiments. Specific statistical tests applied are described in the figure legends. Gene set enrichment analyses (GSEA) were performed in RStudio with the clusterProfiler package (85) using the GSEA function with MsigDB-derived gene sets and Benjamini-Hochberg multiple tests correction. Transcription factor activity inference was performed using the decoupleR package (86). Patient survival data were fit to Kaplan-Meier survival functions using the survival package in TCGAbiolinks and differences in overall survival between groups were determined by log-rank test. CRISPR screen statistical analysis was performed using the MAGeCK Counts pipeline (87) within the Galaxy web server (88).

### Data availability

Gene expression data associated with Figure 1 is deposited in GEO with the accession code GSE320331 and gene expression data associated with Figure 4 is deposited in GEO with the accession code GSE306515. Gene expression data associated with Figure 6 is deposited in GEO with the accession code GSE288530. Values for all data points in graphs, differential gene expression and CRISPR screening scores are reported in the Supporting Data Values file.

## Author Contributions

Colin Sheehan

Conceptualization; Methodology; Investigation; Data Curation; Formal Analysis; Visualization; Writing – Original Draft; Writing – Review & Editing; Project Administration; Supervision

Lyndon Hu (Co-second author)*

Conceptualization; Methodology; Investigation; Data Curation; Formal Analysis; Visualization; Writing – Review & Editing

Guillaume Cognet (Co-second author)*

Conceptualization; Methodology; Investigation; Data Curation; Formal Analysis; Visualization; Writing – Review & Editing

Grace Croley (Co-second author)*

Conceptualization; Methodology; Investigation; Data Curation; Formal Analysis; Visualization; Writing – Review & Editing

Hu, Cognet and Croley contributed equally to this manuscript, order was decided randomly. Rachel Nordgren

Thao Trang Nguyen

Investigation; Formal Analysis

Anika Thomas-Toth

Investigation; Formal Analysis

Darby Agovino

Investigation; Formal Analysis

Patrick B. Jonker

Investigation; Formal Analysis

Mumina Sadullozoda

Investigation; Formal Analysis

Leah M. Ziolkowski

Investigation; Formal Analysis; Writing – Review & Editing

James K. Martin 2^nd^

Investigation; Formal Analysis; Conceptualization

Alica K. Michels

Investigation; Formal Analysis; Writing – Review & Editing

Ranya Dano

Investigation; Formal Analysis; Methodology

Mohammed A. Khan

Investigation; Formal Analysis; Methodology

Rasangi Perera

Investigation; Formal Analysis; Methodology

Isabel Alcazar Investigation

Christopher J. Halbrook

Methodology; Investigation; Project Administration

Kay F. Macleod

Investigation; Writing – Review & Editing

Hardik Shah

Investigation; Formal Analysis; Methodology; Writing – Original Draft; Writing – Review & Editing; Project Administration; Supervision

Christopher R. Weber

Investigation; Methodology; Writing – Review & Editing; Project Administration; Supervision

James L. LaBelle

Investigation; Methodology; Writing – Review & Editing; Project Administration; Supervision

Alexander Muir

Conceptualization; Methodology; Formal Analysis; Writing – Original Draft; Writing – Review & Editing; Project Administration; Supervision

## Funding

This work is the result of NIH funding, in whole or in part, and is subject to the NIH Public Access Policy. Through acceptance of federal funding, the NIH has been given a right to make the work publicly available in PubMed Central.

## Supporting information

Supporting data values

## Acknowledgements

We thank all members of the Muir lab, Jon Coloff, Benjamin Stanger and Yogev Sela for feedback and discussions on this project. We thank Chris Chidley for advice on genetic screening, the University of Chicago Animal Resources Center (RRID:SCR_021806), especially Ani Solanki, for their assistance with animal models of pancreatic cancer and Ernst Lengyel for assistance with tissue sectioning. We thank the University of Chicago Genomics Facility (RRID:SCR_019196), Human Tissue Resource Center (RRID:SCR_019199), Integrated Light Microscopy Core (RRID: SCR_019197), Organoid and Primary Culture Research Core (RRID:SCR_022935) and the Cellular Screening Center (RRID:SCR_017914) for experimental assistance. This work was supported by funding from the Ludwig Center for Metastasis Research, the University of Chicago Comprehensive Cancer Center, the Krull Pancreatic Cancer Award, the National Cancer Institute (R01 CA276461) and the V Foundation (V Scholar Award) provided to A.M. C.J.H. acknowledges support from the National Cancer Institute (R37 CA283575) and the Sky Foundation. C.S.S. was supported by the National Cancer Institute (T32 CA009594 and F31 CA278362). G.C. was supported by the Pancreatic Cancer Action Network (Pancreatic Cancer Action Network Fellowship, funded by the Francois Wallace Monahan Fund). D.A. and P.B.J. were supported by the National Cancer Institute (T32 CA009594).

## Notes

### Competing Interest Statement

The authors have declared no competing interest.

### Summary of Updates

We have added new Figure 1 and Figure 2 to directly measure tumro perfusion, chemotherapeutic accumulation and biochemical target engagement in pancreatic cancers. We additionally provide new genome-wide CRISPR screening data demonstrating that BCL-XL is required for therapy tolerance in pancreatic cancer. Lastly, we show that amino acid stress is a driver of therapy tolerance in pancreatic cancer.

