## Supplemental methods and data for "Tumor nutrient stress gives rise to a drug tolerant cell state in pancreatic cancer"

**Supplemental data**

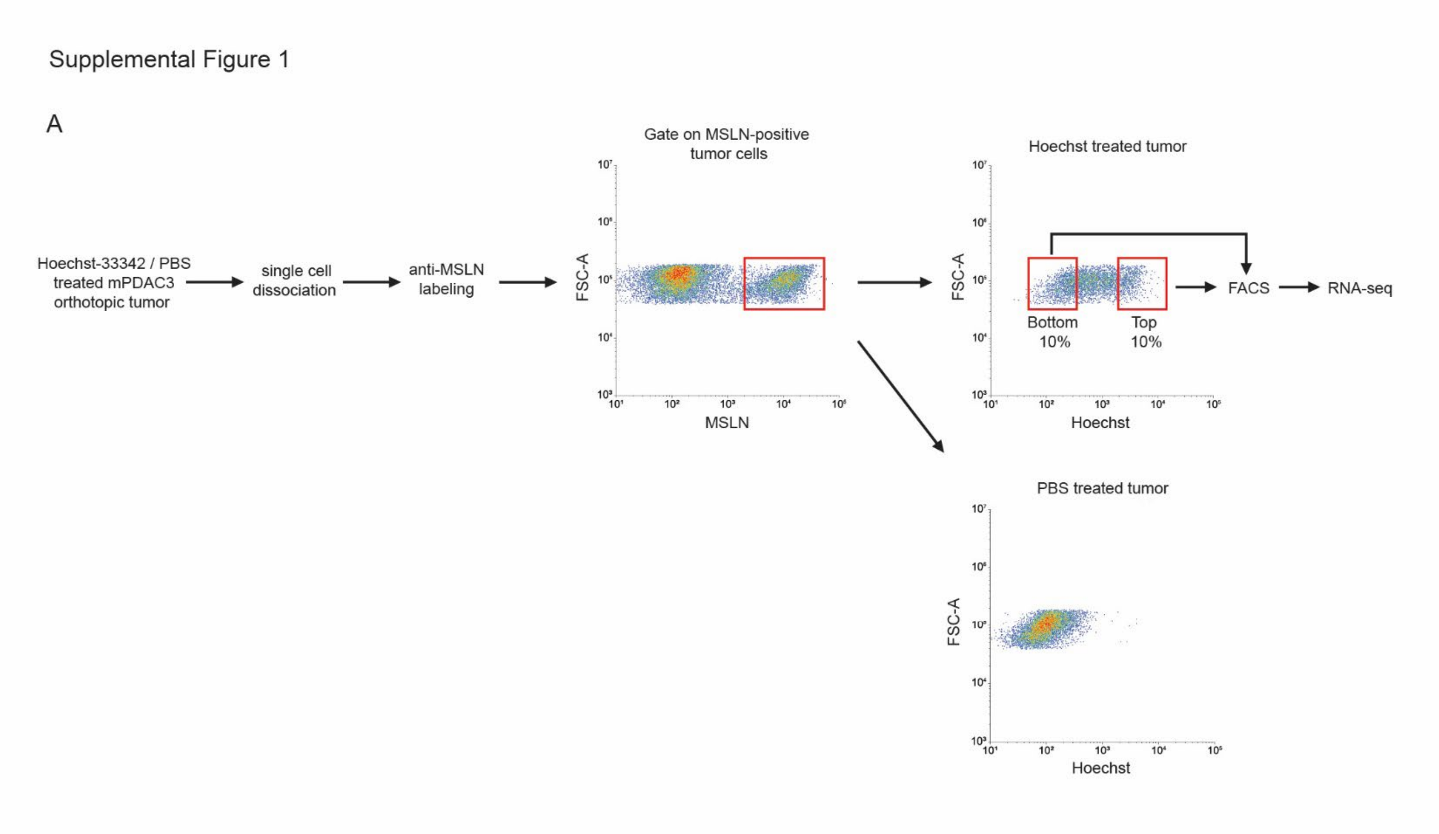


**Supplemental Figure 1: Schema for sorting of well and poorly perfused PDAC cells from tumors. (A)** mPDAC3 cells were identified from orthotopic tumors based on the cell surface marker mesothelin (MSLN). Tumor cells were stratified by local perfusion levels through Hoechst 33342 labeling. The top and bottom 10% Hoechst 33342 labeled cells were collected for transcriptomic analysis.

Supplemental Figure 2

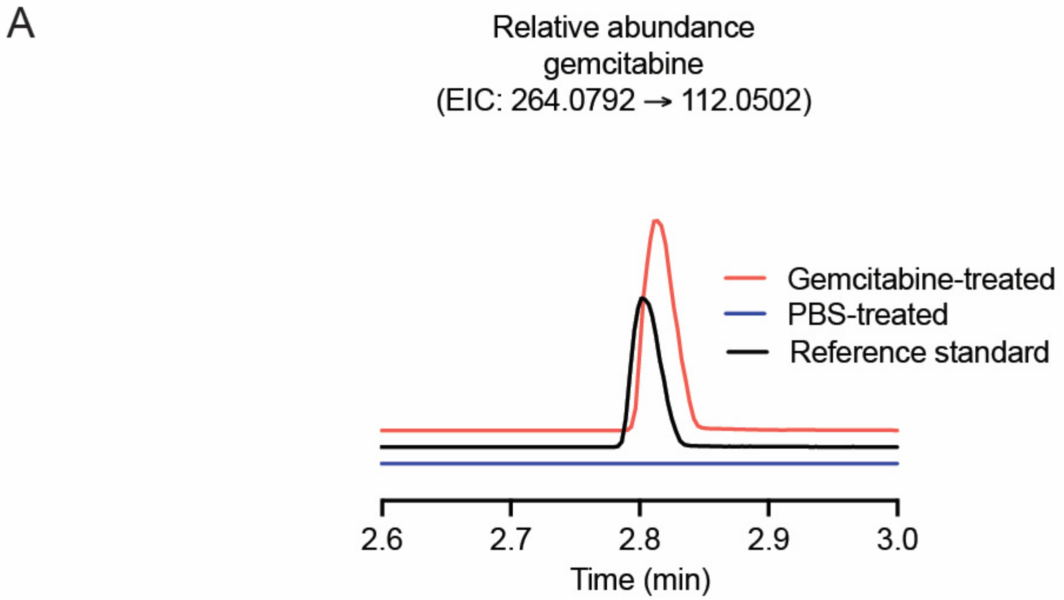


**Supplemental Figure 2: Chromatograms of gemcitabine within biofluids. (A)** Gemcitabine levels were quantified from plasma and tumor interstitial fluid (TIF) samples based on interpolation from an external standard curve. Shown are chromatograms for peak associated with gemcitabine (m/z 264.079 →112.0502) with representative samples from mice treated with and without gemcitabine and a reference standard.

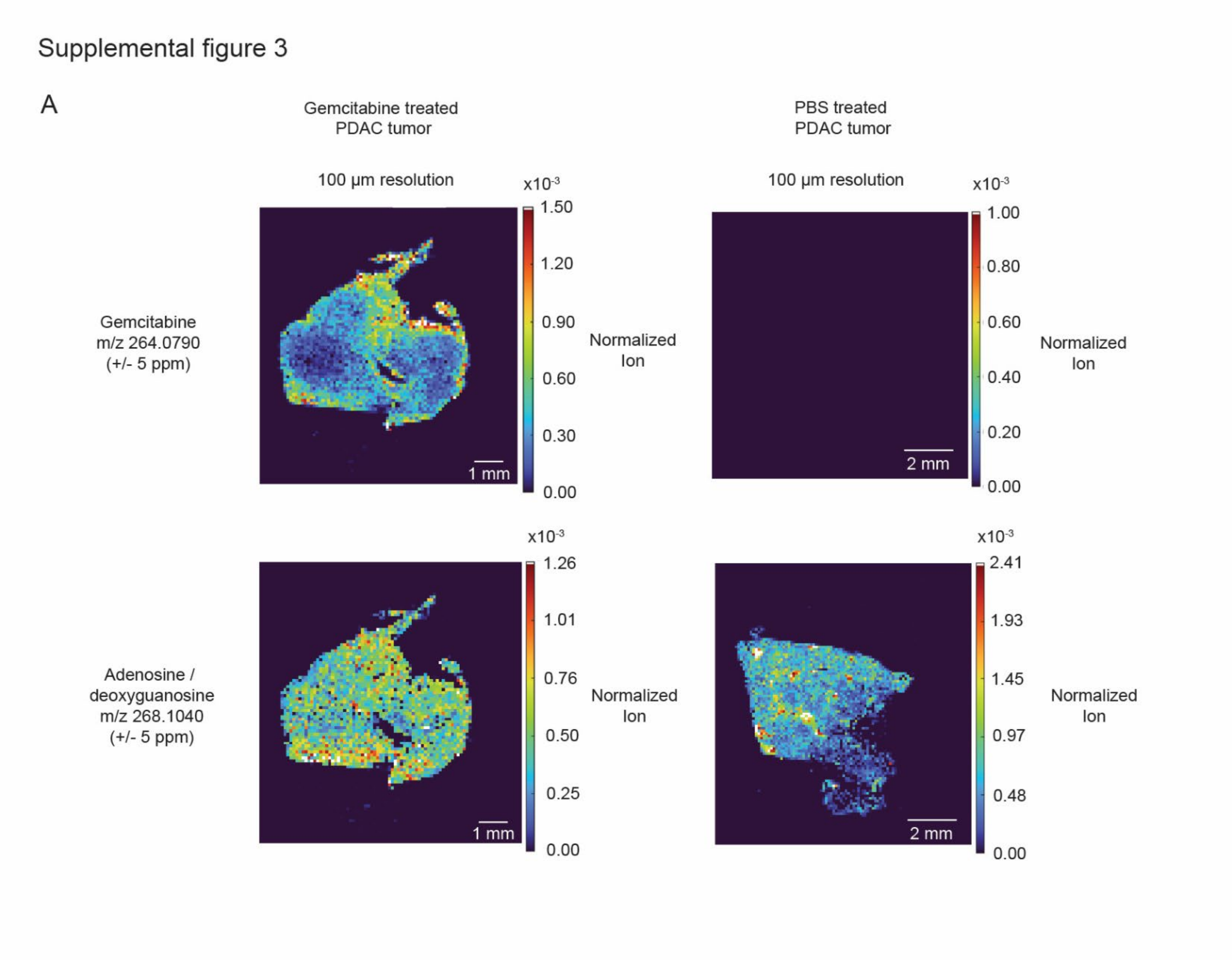


**Supplemental Figure 3: Mass spectrometry imaging identification of gemcitabine in PDAC tumors. (A)** Levels of gemcitabine and the endogenous metabolites adenosine/deoxyguanosine were measured in mPDAC3-RPMI tumors treated with gemcitabine or vehicle (PBS) 1 hour post-treatment by AP-MALDI mass spectrometry imaging. Scale bars correspond to 1 mm for the gemcitabine treated tumor (left) or 2 mm for the PBS treated tumor (right).

Supplemental Figure 4

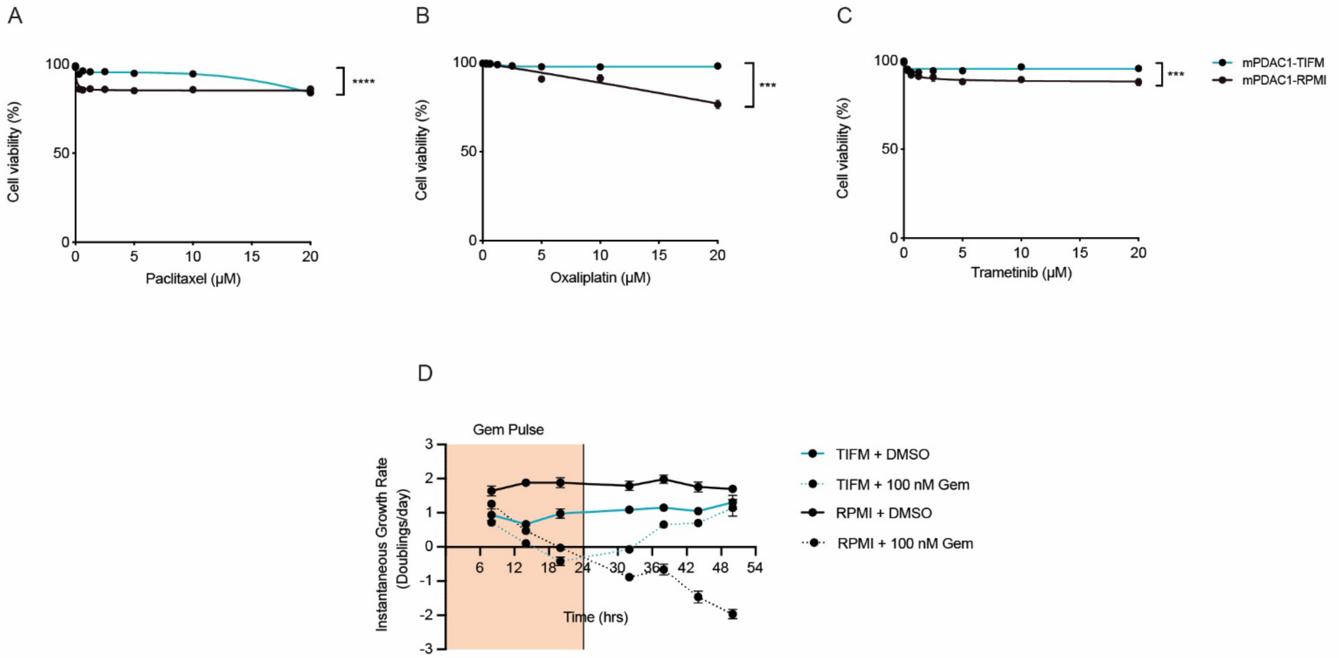


**Supplemental Figure 4: TIFM culture causes PDAC cells to be resistant to common chemo- and targeted therapies. (A-C)** Cell viability of mPDAC1-RPMI and mPDAC1-TIFM cells following treatment with **(A)** paclitaxel, **(B)** oxaliplatin, and **(C)** trametinib at the indicated concentrations for 72 hours (n = 3). **(D)** Growth rates of mPDAC1-RPMI and mPDAC1-TIFM cultures measured every 6 hours over 2 days following exposure to 100 nM gemcitabine for 24 hours (n = 3). For A-C, the area under the curve (AUC) was calculated and unpaired t-test was performed to determine significance in the difference of AUCs for the indicated cultures. *** p≤0.001 and **** p≤0.0001.

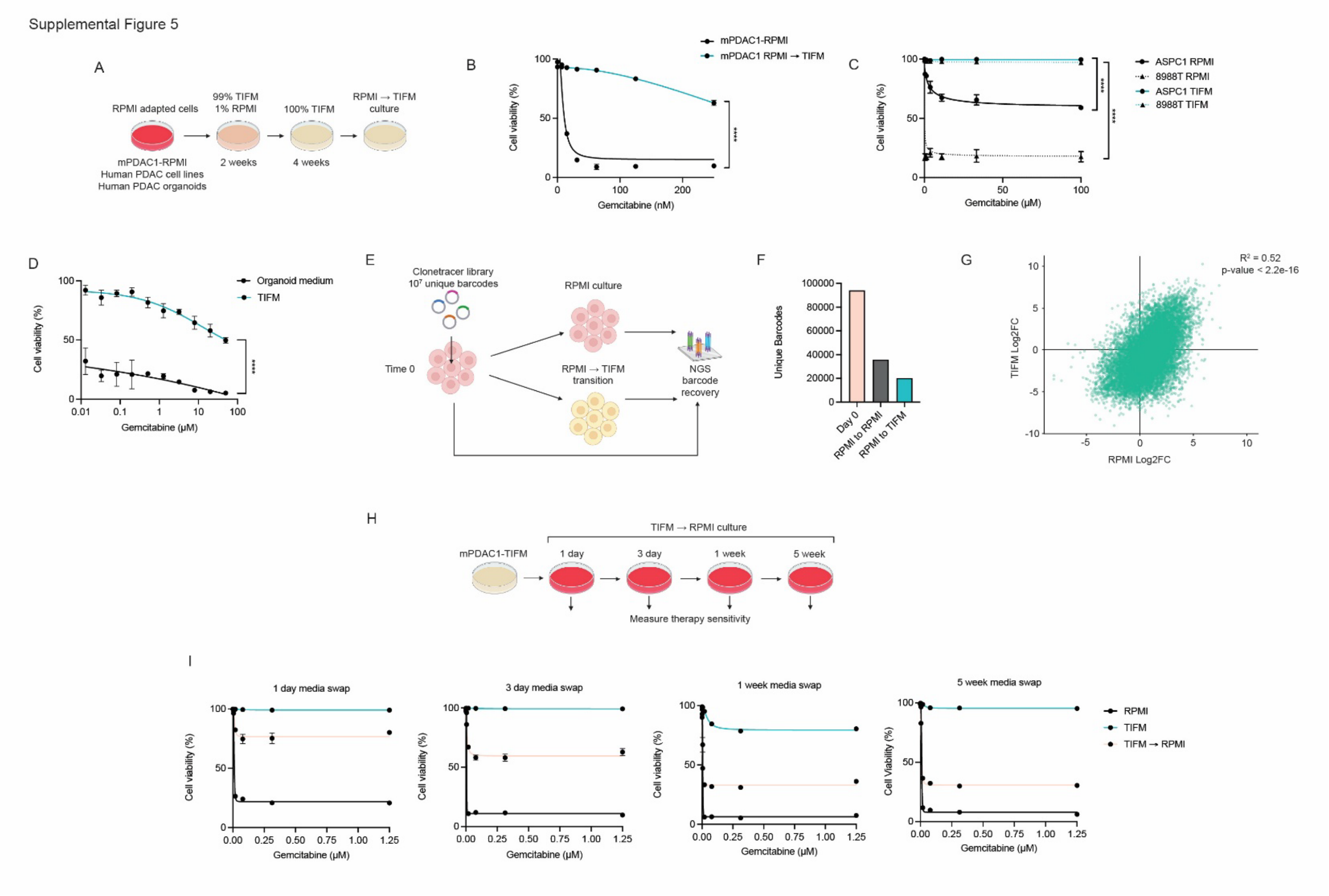


**Supplemental Figure 5: TIFM medium conditions promote a reversible drug resistant cell state which is not clonally selected. (A)** Diagram of the protocol for transitioning mPDAC-RPMI and human PDAC cell lines and organoids to grow in TIFM. We term these mPDAC-RPMI→TIFM cultures. **(B)** Viability of mPDAC1-RPMI and mPDAC1-RPMI→TIFM cultures when treated with the indicated concentration of gemcitabine (n = 3) for 72 hours. **(C)** Viability of indicated human PDAC cell lines cultured in RPMI or TIFM when treated with the indicated concentration of gemcitabine for 72 hours (n = 3)**. (D)** Viability of patient derived organoids measured by CellTiter-Glo assay in standard organoid culture media or TIFM following treatment with the indicated concentration of gemcitabine for 72 hours (n = 3). **(E)** Diagram of the lentiviral barcoding of mPDAC3-RPMI cells prior to transitioning these cells into TIFM or maintenance of the culture in RPMI. **(F)** Barcode recovery of mPDAC3-RPMI cells either maintained in RPMI or transitioned to grow in TIFM. **(G)** Correlation of log2 fold change in barcode abundance between mPDAC-RPMI cells transitioned into TIFM or maintained in RPMI. **(H)** Diagram of experiment transferring mPDAC1-TIFM cultures into RPMI medium (TIFM → RPMI) for various period of time and assessing chemotherapeutic response. **(I)** Cell viability of mPDAC1-TIFM, mPDAC1-RPMI and mPDAC1-TIFM→RPMI cells after the indicated period of time in RPMI and treated with the indicated concentration of gemcitabine for 72 hours (n = 3). For B, C, D, the area under the curve (AUC) was calculated and unpaired t-test was performed to determine significance in the difference of AUCs for the indicated cultures. For G, Spearman's rank correlation test was used to determine significant correlation in change of barcode abundance between culture conditions. **** p≤0.0001.

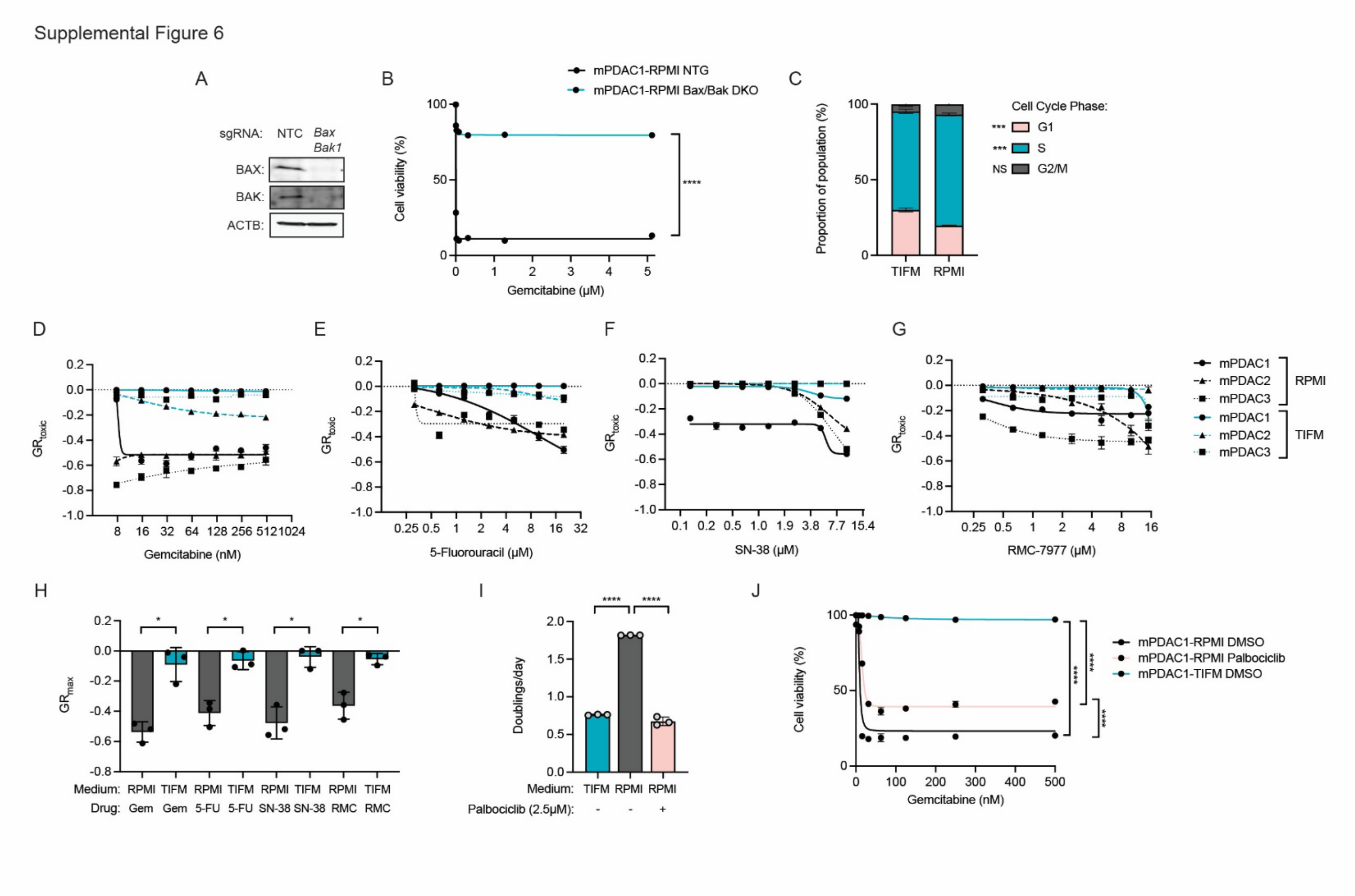


**Supplemental Figure 6: TIFM culture promote a drug resistant cell state not explained by differences in growth rate. (A)** Immunoblot confirmation of *Bax* and *Bak1* double knockout (*Bax/Bak1* DKO) in mPDAC1-RPMI cells. **(B)** Cell viability of mPDAC1-RPMI NTC and *Bax/Bak1* DKO cultures treated with the indicated concentration of gemcitabine for 72 hours (n = 3). **(C)** Percentages of mPDAC1-TIFM cells in G1, S and G2/M phases of the cell cycle following treatment with 100 nM gemcitabine for the indicated time periods (n = 3). **(D-G)** $GR_{toxic}$ estimates for the indicated mPDAC-TIFM and mPDAC-RPMI cell line following treatment with the indicated concentration of **(D)** gemcitabine (n = 3), **(E)** 5-fluorouracil (n = 3), **(F)** SN-38 (n = 3), and **(G)** RMC-7977 (n = 3) for 72 hours. **(H)** $GR_{max}$ estimates for the indicated mPDAC cell lines, culture condition and drug treatment (n = 3). **(I)** Growth rates of vehicle treated mPDAC1-TIFM and mPDAC1-RPMI cultures and mPDAC1-RPMI cultures treated with 2.5 µM palbociclib (n = 3). **(J)** Viability of mPDAC1-TIFM and mPDAC1-RPMI cultures and mPDAC1-RPMI cells treated with 2.5 µM palbociclib cultures following treatment with the indicated concentration of gemcitabine for 72 hours (n = 3). For C, H, I, statistically significant differences were determined by two sample T-test. For B, J, the area under the curve (AUC) was calculated and unpaired t-test was performed to determine significance in the difference of AUCs for the indicated cultures. * p≤0.05 *** p≤0.001 and **** p≤0.0001.

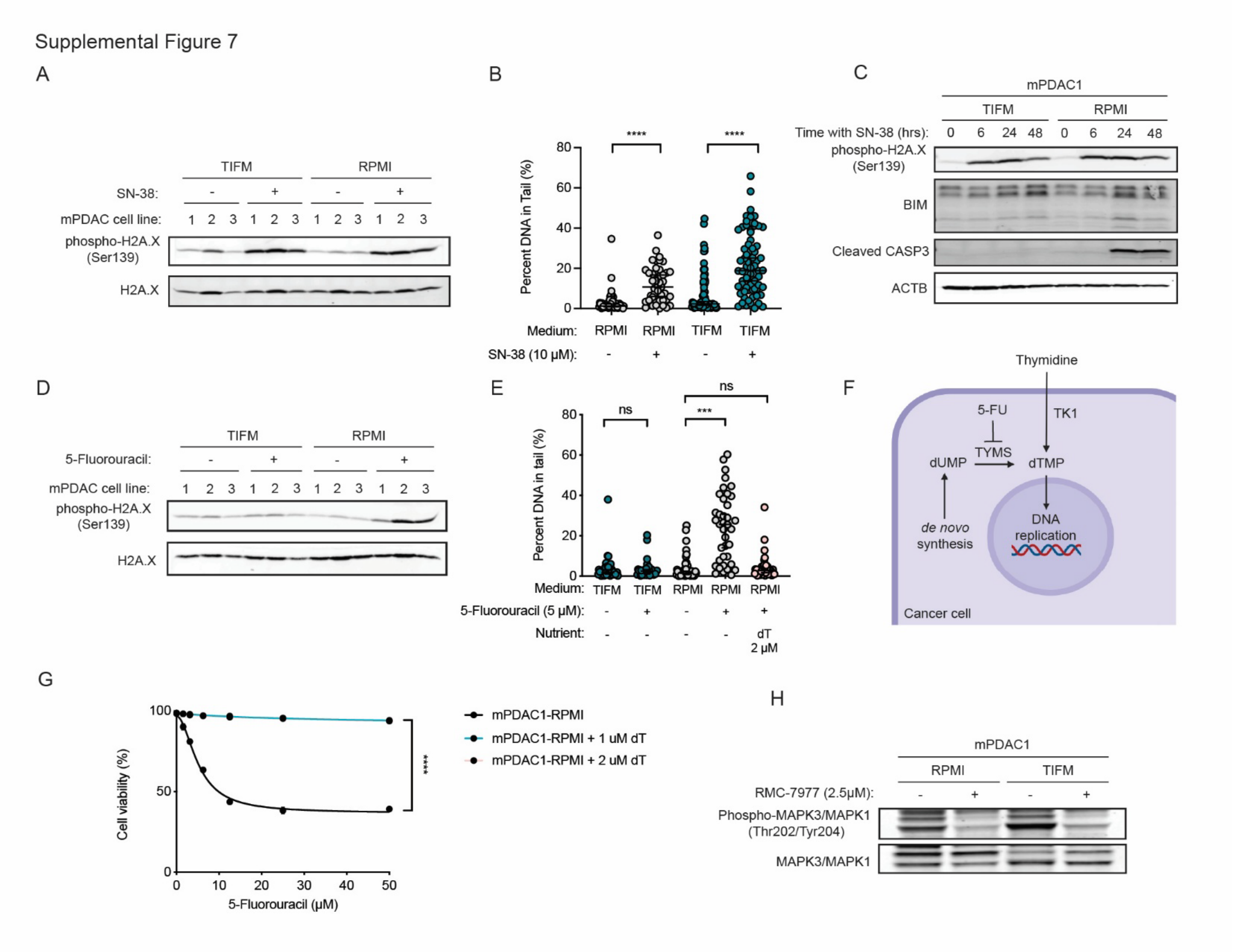


**Supplemental Figure 7: With the exception of 5-fluorouracil, TIFM culture does not interfere with the biochemical activity of therapeutic agents. (A)** Immunoblots for γH2AX and H2AX in the indicated mPDAC cell line and culture condition following treatment with 10 μM SN-38 for 18 hours. **(B)** Alkaline comet assay of mPDAC1-RPMI and mPDAC1-TIFM cells treated with 10 μM SN-38 (RPMI n = 45, TIFM n = 70) or vehicle (RPMI n = 55, TIFM n = 70) for 18 hours. **(C)** Immunoblots for γH2AX, BIM, and cleaved caspase-3 in mPDAC1-RPMI and mPDAC1-TIFM cells treated with 10 μM SN-38 over time at the indicated timepoints. **(D)** Immunoblots for γH2AX and H2AX in the indicated mPDAC cell line and culture condition following treatment with 5 μM 5-FU for 18 hours. **(E)** Alkaline comet assay of mPDAC1-RPMI and mPDAC1-TIFM cells treated with 5 μM 5-FU (RPMI n = 38, TIFM n = 34, RPMI + thymidine n = 46) or vehicle (RPMI n = 38, TIFM n = 50) for 18 hours. **(F)** Diagram of metabolic routes for dTMP synthesis. **(G)** Cell viability of mPDAC1-RPMI cells following treatment with indicated concentrations of 5-FU in the presence or absence of the indicated concentrations of thymidine after 72 hours (n = 3). **(H)** Immunoblots for phospho-ERK and ERK in mPDAC1-RPMI and mPDAC1-TIFM cells treated with 2.5 μM RMC-7977 or vehicle for 24 hours. For B, E, statistically significant differences were determined by two-sample T-test. For G, the area under the curve (AUC) was calculated and unpaired t-test was performed to determine significance in the difference of AUCs for the indicated cultures. *** p≤0.001 and **** p≤0.0001.

### Supplemental methods

#### *Tumor growth studies*

Orthotopic allograft tumors were established with mPDAC3-RPMI cells as described above. For tumor growth studies, gemcitabine treatment was initiated 2 weeks after initial tumor implantation. Gemcitabine hydrochloride (USP, 1288463) was resuspended in PBS, sterilized by syringe filtration, and dosed at 120 mg/kg via intraperitoneal injection. Tumor-bearing mice received gemcitabine or PBS injections twice weekly for 3 weeks. Mice were euthanized 24 hours after the final treatment with gemcitabine and tumor weights were measured.

#### *Tumor vascularization and perfusion studies*

Orthotopic allograft tumors were established with mPDAC3-RPMI cells as described above. After 3 weeks of tumor growth, mice were injected via tail vein with 10 mg/kg Texas red-conjugated *Lycopersicon esculentum* lectin (LEL) (Bioworld #21761032) resuspended in PBS and incubated for 25 minutes. Following 25 minute incubation period, mice were additionally injected via tail vein with 20 mg/kg Hoechst 33342 (Sigma B2261) resuspended in PBS and incubated for 5 minutes. Following the 5 minute incubation period, mice were euthanized via cervical dislocation and primary tumors were rapidly dissected and placed in pre-chilled 4% paraformaldehyde and stored at 4 °C for 5 hours. Fixed tumors were then transferred to pre-chilled 15% sucrose solution in PBS at 4 °C until equilibration, then transferred to pre-chilled 30% sucrose solution in PBS at 4 °C overnight. Tumor specimens were then embedded in OCT reagent and rapidly frozen. 10 µm cryosections were generated with the assistance of the UChicago Human Tissue Resource Center (HTRC) (RRID:SCR_019199) and stored at -80 °C until the time of imaging and analysis as described above.

#### *Cyclic immunofluorescence studies (t-CyCIF)*

Methods for cyclic immunofluorescence were adapted from Lin et al., 2018 (13). Tumor specimens were initially imaged to measure perfusion markers (perfused Hoechst 33342 and or Texas red conjugated LEL without significant manipulation of the tissue. Frozen tissue sections were thawed, rinsed twice with 1x PBS solution, and coverslips were mounted with 1x PBS solution with 10% glycerol. Following the initial round of imaging, coverslips were removed by submerging in 1x PBS solution until they naturally fell off. Slides were then subjected to antigen retrieval in citrate buffer (10mM citrate, 0.05% tween-20, pH 6.0) via pressure cooker (Instant Pot 113-0059-01) (110 °C for 5 minutes). Slides were then rinsed once in PBS-T (137mM NaCl, 2.7mM KCl, 10mM $Na_2HPO_4$, 1.8mM $KH_2PO_4$, 0.1% Tween-20 (w/v), pH 7.4) buffer and subjected to an initial round of destaining. Slides were submerged in fluorophore bleaching solution (20 mM NaOH, 4.5% $H_2O_2$, in PBS) and placed directly under a high intensity LED light panel for 1 hour. Slides were then rinsed twice in PBS-T buffer and blocked for 3 hours at 4 °C with Intercept blocking buffer (LICOR 92770001). All staining was performed using conjugated primary antibodies (PE-conjugated phosho-H2A.X (Ser139) (Cell Signaling 9718), AF488-conjugated cleaved-caspase-3 (Asp175) (Cell Signaling 9669S), AF647-conjugated Arginase-1 (Cell Signaling 43279S), AF555-conjugated cleaved-Parp (Asp214) (Cell Signaling 6894S), CD31 (Cell Signaling 77699S), AF488-conjugated F4/80 (Biorad MCA497A488T)) diluted at 1:100 dilution in a 1:1 mix of intercept blocking buffer with PBS-T, in the presence of 300nM Hoechst 33342 as a nuclear marker. Slides were then incubated at 4 °C overnight in a humidified chamber. In cases where a directly conjugated primary antibody was not available, antibody conjugation was performed prior to staining using the proteintech FlexAble linker kit (Proteintech KFA503). The following day, slides were washed 3x with PBS-T buffer, mounted with coverslips, imaged, bleached, and restained as described. All rounds of imaging were performed using a Leica SP8 Laser Scanning Confocal Microscope. Image tiling for large tissue sections was performed using the Leica software and exported as TIF files. Image registration was guided using the total nuclear Hoechst 33342 marker for each image set, using the MultiStackRegistration ImageJ plugin with a rigid body algorithm. Background subtraction for each image was performed based on the median pixel intensity of unstained autofluorescence control slides collected during each round of imaging. Registered images were then merged into hyperstacks and converted to OME TIFF files for further analysis using QuPath software (14).

*Quantification of lectin in tumor vessels and quantification of perfused Hoechst 33342 around tumor blood vessels*

Regions of interest covering the entire tumor section were manually drawn for each tumor section. A pixel classifier was trained using CD31 fluorescence intensity and manual annotation of vessels within representative images. This classifier was then loaded to our experimental image set to generate vessel objects across the sample, resulting in at least 1000 objects per specimen. Local intensity features were quantified for LEL fluorescence and perfused Hoechst 33342 fluorescence and added to each vessel object. For perfused hoechst measurements, a preferred pixel size of 25 μm was used to include vessel adjacent cells within local intensity quantification. Spearman's correlation on the LEL and perfused Hoechst 33342 measurements for each vessel was then performed to determine whether there was a relationship between the local intensity of these perfusion markers.

*Quantification of perfusion markers in normal pancreas vs. allograft orthotopic tumors.*

Regions of interest covering the entire tumor section were manually drawn for each tumor section. CD31+ blood vessels were identified using the pixel classifier described above. Local fluorescence intensity measurements for LEL were added to each vessel. The median of all vessel measurements was then determined for each tissue specimen. For perfused Hoechst 33342 measurements, individual cells within each tissue specimen were identified using the cell detection feature guided by nuclear staining. For each cell detected, the median perfused Hoechst 33342 intensity was determined. The median value of all cell measurements was determined for each tissue specimen.

*Quantification of correlation between local perfusion rate and gemcitabine level*

Images for gemcitabine levels were collected via AP-MALDI mass spectrometry imaging (MSI) as described below, and images for local perfusion levels were collected via confocal fluorescent microscopy with Hoechst 33342 using parallel tumor sections. In brief, total ion current (TIC) normalized intensity images for the gemcitabine main fragment ion (m/z 112.0502±2.5 ppm) were exported as grayscale RGB images from MSiReader software. Simultaneously, perfused Hoechst 33342 and total nuclear Hoechst 33342 images were acquired and registered as described above. Fluorescent microscopy images were scaled to match MSI images and were manually registered to the MSI images within ImageJ. Registered images were then all converted to 8-bit images and merged as separate channels for further analysis within QuPath software. Regions of interest covering the entire tumor section were manually drawn for each sample. Tumor ROIs were then segmented into 100 pixel SLIC superpixel regions, covering approximately 100 ROIs per tumor section. Local intensity features were added to superpixel annotations composed of the median intensity of the perfused Hoechst channel and the median intensity of the gemcitabine channel. Spearman's rank coefficient correlation method was utilized to determine the correlation between Hoechst perfusion and gemcitabine levels for each tumor section.

*Total quantification of γH2AX and cleaved-caspase-3 in tumor sections*

Regions of interest covering the entire tumor section were manually drawn for each tumor section. Positively labeled cells for phospho-H2AX or cleaved caspase-3 were determined using the positive cell detection function in QuPath using the nuclear label to facilitate total cell detection. A thresholding cutoff for positive cells was determined based on the mean pixel intensity for a sample of positive cells identified from a representative tumor image.

*Quantification of DNA damage rate as a function of local perfusion*

Positively labeled cells for phospho-H2AX within each tumor section were determined as described above. For each cell, the fluorescence intensity in the Hoechst 33342 channel was determined and binned into percentiles based on median fluorescence intensity of the total tumor cell population. The proportion of phospho-H2AX-positive cells was then calculated for each population bin. Linear regression was then performed to determine whether there was a relationship between rates of phospho-H2AX positivity and Hoechst 33342 labeling levels.

_Quantification of arginase-1 expression as a function of local perfusion_

Regions of interest covering the entire tumor section were manually drawn for each tumor section. Cells within each tissue specimen were identified using the cell detection feature guided by nuclear staining. For each cell detected, the median perfused Hoechst 33342 intensity and median arginase-1 intensity was determined. Linear regression was then performed to determine whether there was a relationship between Hoechst 33342 labeling and arginase-1 expression level.

***Transcriptomic Studies***

_Well and poorly perfused PDAC cells_

Orthotopic allograft tumors were established with mPDAC3-RPMI cells as described above. Mice were euthanized 3 weeks after initial tumor implantation for RNA-seq analysis. Just prior to euthanasia, mice were treated with 20 mg/kg of Hoechst 33342 33342 dye or PBS via tail vein injection and euthanized by cervical dislocation after 5 minutes. Tumors were then rapidly dissected and dissociated as described previously (1). Single cell tumor suspensions were then stained for mesothelin expression to identify PDAC cells. Tumor suspensions were initially incubated with 10% normal mouse serum (Thermo, 10410) in PBS containing 1% BSA for 15 minutes at 4 °C to block Fc receptors. Samples were then stained with rat anti-mesothelin (mouse) (MBL, D233-3) antibody at 1:100 dilution in 10% normal mouse serum in PBS containing 1% BSA for 30 minutes at 4 °C. Samples were washed twice with PBS containing 1% BSA and then stained with APC-conjugated goat anti-rat IgG (Thermo, A10540) secondary antibody at 1:200 dilution in 10% normal mouse serum in PBS containing 1% BSA for 20 minutes at 4 °C. Samples were then washed twice with PBS containing 1% BSA and subjected to FACS. Cancer cells were initially gated based on positive mesothelin staining, and high and low perfused Hoechst 33342 label was identified based on the top and bottom 15% of the cancer cell population. High and Low perfused populations were then sorted directly into RLT lysis buffer for RNA extraction, with a minimum of 15,000 cells sorted per sample. RNA was extracted using the RNeasy Micro Kit (Qiagen, 74004). RNA quality and quantity were then assessed using the 2100 Bioanalyzer System (Agilent).

_Gemcitabine response in mPDAC-TIFM vs. mPDAC-RPMI cells_

mPDAC1 RPMI and TIFM cultured cells were plated at 50,000 cells per well in 6 cm dishes overnight. The following day, cells were treated with vehicle or 125 nM gemcitabine. After 24 hours of treatment, RNA was extracted from treated cells using the RNeasy Micro Kit (Qiagen, 74004). RNA quality and quantity were then assessed using the 2100 Bioanalyzer System (Agilent).

_Library preparation and sequencing_

Strand-specific RNA-SEQ libraries were prepared using the TruSEQ mRNA RNA-SEQ library protocol (Illumina). Library quality and quantity were assessed using the Agilent bio-analyzer and libraries were sequenced using an Illumina NovaSEQ6000 platform at the University of Chicago Functional Genomics Facility (RRID:SCR_019196).

_RNA Sequencing data analysis_

Data processing and analysis were done using the R-based Galaxy platform (https://usegalaxy.org/). Quality control was performed prior and after concatenation of the raw data with the MultiQC and FastQC tools, respectively. All samples passed the quality check with most showing ~20% sequence duplication, sequence alignment greater or equal to 80%, below and below 50% GC coverage. Samples were aligned and counts were generated using HISAT2 (Galaxy Version 2.2.1+galaxy0, NCBI genome build GRCm38/mm10) and featureCounts (Galaxy Version 2.0.1+galaxy1), respectively. Differential expression analyses were performed with limma (Galaxy Version 3.48.0+galaxy1). Gene set enrichment analyses (GSEA) were performed in RStudio with the clusterProfiler package (15) using the GSEA function with C2 curated gene sets from MsigDB and Benjamini-Hochberg multiple tests correction. Altered gene set enrichments were determined by initially separating gene set enrichments into positive and negatively enriched gene sets, sorting by the lowest adjusted p-value, and then sorting by the highest normalized enrichment score. For the krige amino acid deprivation gene set enrichment in mPDAC1-TIFM vs. mPDAC1-TIFM + R as well as the hallmark apoptosis gene set enrichment in gemcitabine vs. DMSO treated PDAC cells, GSEA analysis was performed as described above; however, testing only a single gene set and therefore, multiple tests correction was not applied.

*Correlation of differentially expressed genes in mPDAC1-TIFM vs. mPDAC1-RPMI cells with Gemcitabine treatment*

Differential expression analysis for genes altered with gemcitabine treatment in mPDAC1-RPMI and mPDAC1-TIFM cells was performed as described above. Differentially expressed genes were defined based on a log2 fold change greater than 1 or less than -1 with an adjusted p-value less than 0.05. Differentially expressed genes between RPMI and TIFM conditions were plotted as a fold-change vs. fold-change plot and spearman's rank correlation coefficient method was applied to determine the correlation of gemcitabine-altered genes between RPMI and TIFM conditions.

*Transcription factor activity inference analysis*

Transcription factor activity inference for mPDAC1-RPMI and mPDAC-TIFM differential gene expression datasets with gemcitabine treatment was performed using the decoupleR package (16). Analysis was performed using a univariate linear model (ULM) for predicted TF activity based on known TF-gene interactions and observed differential gene expression with the run_ulm() function. The murine CollecTRI gene regulatory network was used as input for the known TF-gene interactions. Transcription factors with predicted enhanced activity were determined based on the highest ULM enrichment score.

*Poor perfusion gene enrichment in chemotherapy unresponsive tumors*

A poor perfusion gene set was generated based on differential gene expression of low vs. high Hoechst 33342 labeled cells, which included genes with $\log_2$ fold change > 1.0 and an adjusted p-value < 0.01. Transcriptomic data in the form of STAR count files for primary tumor specimens from the full TCGA pancreatic cancer cohort (TCGA-PAAD) were downloaded in R using the TCGA biolinks package. Primary treatment response data was downloaded from the TCGA Broad GDAC Firehose server (https://gdac.broadinstitute.org/). Patients were filtered for those with pancreatic adenocarcinoma ductal histological subtype, stage IIa/IIb pathological staging, and receiving chemotherapy within their initial treatment. RNA expression count files were transformed using vst normalization with the DESeq2 package. Gene set variation analysis (GSVA) (17) was used to generate pathway enrichment scores for each specimen using the poor perfusion gene set with the GSVA package. Patients were then categorically binned based on measure of response within clinical metadata into either responders ("complete response" or "partial response") or non-responders ("stable disease" or "progressive disease"). Differences in enrichment of poor perfusion genes between responder and non-responder tumors were then evaluated by two sample t-test.

*Poor perfusion & Krige amino acid deprivation gene set enrichment effects on overall survival in PDAC*

A poor perfusion gene set was generated based on differential gene expression of low vs. high Hoechst 33342 labeled cells, which included genes with $log_2$ fold change > 1.0 and an adjusted p-value < 0.01. The krige amino acid deprivation gene set was downloaded from the murine C2 gene sets with the msigdbr package. Transcriptomic data in the form of STAR count files for tumor specimens from the full TCGA pancreatic cancer cohort (TCGA-PAAD) were downloaded in R using the TCGA biolinks package. RNA expression count files were transformed using vst normalization with the DESeq2 package. Gene set variation analysis (GSVA) (17) was used to generate pathway enrichment scores for each specimen using the poor perfusion gene set or the krige amino acid deprivation gene set with the GSVA package. Patients were then stratified into high and low enrichment of poor perfusion genes or krige amino acid deprivation genes based on the median GSVA enrichment score across all samples. Patient survival data was downloaded using TCGA biolinks and the high/low enrichment groups were fitted to Kaplan-Meier survival functions using the survival package. Differences in overall survival between groups were determined by log-rank test.

***Measurements of gemcitabine in TIF***

*TIF extraction from gemcitabine treated mice*

Orthotopic allograft tumor bearing mice were treated with a single intraperitoneal injection of gemcitabine at 120 mg/kg, or PBS, after 3 weeks following the initial tumor implantation. Following gemcitabine treatment, mice euthanized after 1 hour. Just prior to euthanasia by cervical dislocation, blood samples were collected via cheek bleed in EDTA-coated collection tubes and plasma was isolated by centrifugation at 1000 G for 15 minutes at 4 °C. Primary tumors were rapidly dissected, rinsed with sterile saline, and TIF was isolated by centrifugation at 400 G for 10 minutes at 4 °C. TIF and plasma samples were then flash frozen in LN2 and stored at -80 °C until the time of analysis via LC/MS.

*Gemcitabine quantification by LC/MS*

Gemcitabine hydrochloride reference standard was purchased from United States Pharmacopeia (USP 1288463). All the solvents used were LC-MS grade and were purchased from Thermo Fisher Scientific. 2 µL of plasma/TIF was mixed with 30 µL of ice-cold MeOH containing 0.1% formic acid, vortexed, incubated at 4°C for 20 minutes, centrifuged at 18,000 g for 20 mins at 4°C. 10 µL of supernatant was mixed with 90 µL of $H_2O$ containing 0.1% formic acid, vortexed, incubated, centrifuged and the supernatant was transferred to an autosampler vial for LC-MS analysis. Stock solutions of gemcitabine (1mg/ml) were prepared by dissolving the accurately weighed reference compound in water. Calibration standards with the following ranges: (0.195, 0.391, 0.781, 1.593, 3.125, 6.25, 12.5, 25, 50 and 100 µg/ml) were prepared by diluting the stock solution in human albumin (4% w/V in PBS). Chromatography separation was performed using the Thermo Scientific Vanquish Horizon UHPLC system and Waters™ XSelect Premier HSS T3 (100Å, 2.5 µm, 2.1 X 100 mm) and detected using high-resolution Orbitrap ID-X Tribrid mass spectrometer (Thermo Scientific) with a H-ESI probe operating in positive mode. The mobile phase A (MPA) was 0.1% formic acid in water MPB was 90/10 IPA/water. The column temperature, injection volume, and flow rate were 40°C, 2 µL, and 0.4 mL/min, respectively. The chromatographic gradient was 0 min: 0% B, 1 min: 0% B, 3 min: 30% B, 4 min: 100% B, 8 min: 100% B, 8.1 min: 0% B and 12 min: 0% B. MS parameters were as follows: Acquisition at 30K resolution, spray voltage:3600V for positive ionization mode, sheath gas: 40, auxiliary gas: 8, sweep gas: 1, ion transfer tube temperature: 250°C, vaporizer temperature: 350 °C,maximum injection time of 54 ms and normalized HCD(%):30. Gemcitabine identification was done by matching the retention time and fragmentation of the precursor ion at m/z 264.079 →112.0502 and quantify using the external calibration curve approach. Data acquisition was done using the Xcalibur software (Thermo Scientific) and quantitative analysis was performed using TraceFinder 5.1 software (Thermo Scientific).

***Mass spectrometry imaging of gemcitabine in tumors***

*Tissue collection and MALDI data acquisition*

Orthotopic allograft tumor bearing mice were treated with a single intraperitoneal injection of gemcitabine at 120 mg/kg, or PBS, after 3 weeks following the initial tumor implantation. Following gemcitabine treatment, mice euthanized after 1 hour. Just prior to euthanasia by cervical dislocation, mice were injected via tail vein with 20mg/kg Hoechst 33342 and incubated for 5 minutes. Primary tumors were rapidly dissected, rinsed with sterile saline, and flash frozen in LN2 and stored at -80 °C. Frozen tumor tissues were sectioned into 10-μm-thick sections via cryostat (Leica CM1850UV Cryostat) maintained at -20°C. These sections were transferred onto Indium Tin Oxide-coated glass slides (Delta Technologies, Limited, CO, USA) and stored at -80$^0$C until the analysis. Before matrix deposition slides were thawed to room temperature in a desiccator for 30 min. For imaging, α-Cyano-4-hydroxycinnamic acid (CHCA) matrix (10 mg/mL in 70% ACN in 0.01% TFA) was applied in 4 passes using the HTX-M3 -sprayer (HTX-Technologies, Chapel Hill, NC, USA) with the following parameters; temperature, 75 °C; flow rate, 120 μL/min; nozzle velocity, 1200 mm/min; track spacing, 3.0 mm; and $N_2$ pressure, set to 10 psi. After matrix deposition, slides were dried in a vacuum desiccator for 30 min.

All imaging experiments were performed on the AP-MALDI ion source (MassTech, Colombia, Maryland, USA) coupled to the Orbitrap ID-X Tribrid mass spectrometer (Thermo Fisher Scientific, San Jose, CA, USA). For gemcitabine detection, data acquisition was performed in positive ion mode with a targeted m/z of 264.0790 and identifying fragment at m/z 112.0502, HCD 30%, resolution-30K. For adenosine/deoxyguanosine detection, data were acquired in positive mode (m/z 200 – 310), 60K resolution. The MS parameters were: spray voltage:2000V, ion transfer tube temperature: 275°C, with a diode-pumped solid-state laser operating at 1.5% laser energy, 3000 Hz frequency. Spectra were acquired over each tissue section at 100μm spatial resolution.

*MALDI Data processing and annotation*

Raw MSI data were processed in SViewer 2.1.0 and exported in .imzML mode to MSiReader v 3.14. The gemcitabine images were created by plotting the intensity of the main fragment ion (m/z 112.0502±2.5 ppm) as a function of position over the tissues. For intensity of the adenosine/deoxyguanosine, [M+H] ion (m/z 268.104) was plotted as a function of position with a mass tolerance of 5 ppm.

***Time course immunofluorescence analysis of tumors after gemcitabine treatment***

For immunofluorescence studies, mice were treated with a single intraperitoneal injection of gemcitabine at 120 mg/kg after 3 weeks following the initial tumor implantation. After gemcitabine treatment, mice were euthanized at the following timepoints: 0 hour (untreated), 24 hours, and 48 hours. Just prior to euthanasia, mice were treated with 20 mg/kg of Hoechst 33342 dye via tail vein injection and euthanized by cervical dislocation after a 5 minute incubation period. Primary tumors were then rapidly dissected and placed in pre-chilled 4% paraformaldehyde and stored at 4 °C for 5 hours. Fixed tumors were then transferred to pre-chilled 15% sucrose solution in PBS at 4 °C until equilibration, then transferred to pre-chilled 30% sucrose solution in PBS at 4 °C overnight. Tumor specimens were then embedded in OCT reagent and rapidly frozen. 10 μm cryosections were generated with the assistance of the UChicago Human Tissue Resource Center (HTRC) and stored at -80 °C until the time of imaging and analysis as described above.

***Production of Nuclight expressing cancer cell lines***

Murine and human PDAC cell lines were engineered to stably express a nuclear-restricted red fluorescent protein (Nuclight; Sartorius, BA-04887) to facilitate live cell counting using the manufacturer's protocol. For mPDAC1-3 cell lines, CRISPR/Cas9 gene editing was utilized to knockout YFP expression. sgRNAs targeting

YFP were designed and protospacer sequences targeting YFP were purchased as single strand oligonucleotides from IDT using standard desalting purification. Oligonucleotides were then cloned into the lentiCRISPRv2-Blast vector (Addgene, 83480) using previously published methods (5). mPDAC1-3 cells were lentivirally transduced with the resulting vectors as previously described (1). Transduced cells were selected over the course of 5 days with treatment of 5 μg/ml blasticidin and sorted using a BD FACSAria II Cell Sorter to isolate a polyclonal YFP-negative and Nuclight-positive mPDAC culture.

***IncuCyte cell viability and proliferation assays***

Unless otherwise indicated, all chemotherapy sensitivity assays were performed as follows. Cells were plated in 96-well tissue culture plates at a density of 1500 cells/well and allowed to attach overnight. The following day, the media was removed from each well and replaced with fresh media containing Cytotox Green dye (Sartorius, 4633) at 50nM and the indicated concentration of compounds or vehicle. Plates were regularly scanned using a Sartorius/Essen BioScience IncuCyte S3 Instrument to monitor changes in cell viability and cell number over time. Unless otherwise indicated, all data shown reflects cell viability at 72 hours of the relevant treatment. Cellular segmentation and object quantification from the red and green fluorescence channels was performed using the IncuCyte analysis software. Cell viability was quantified based on the proportion of red (Nuclight, viable) and green (Cytotox, dead) objects identified across the entire well, using the following formula: cell viability (%) = $(\frac{\#\, red\ objects}{\#\, red\ objects + \#green\ objects}) * 100$.

In some experiments, cell proliferation rates were determined in response to drug treatments. Here, cell proliferation rates were calculated as the number of population doublings per day, based on the change in the number of red objects from an initial timepoint and a final timepoint. These values were calculated according to the following population growth formula, $doublings\ per\ day = \left(\frac{1}{\#\ days}\right) * log_2(\frac{final\ cell\ count}{initial\ cell\ count})$.

***Normalized growth rate inhibition (GR) analysis***

$GR_{toxic}$ estimates were determined for mPDAC1 cells treated with a titration of targeted and chemotherapy drugs using initial and final live/dead cell counts determined by IncuCyte-based live cell imaging experiments shown in Fig. 2C-F as previously described (8) using the GRcalculator website, (grcalculator.org). $GR_{max}$ estimates were similarly determined.

***Assessment of chemotherapeutic impact on patient-derived organoids***

To test the impact of chemotherapeutics on patient-derived organoids derived in TIFM, TIFM-derived organoids were replated into two 35 uL UltiMatrix domes in a pre-warmed 24-well plate and cultured in one of two media: advanced DMEM/F12 or TIFM basal medium. After three days, organoids were replated as 10 μL UltiMatrix domes into a pre-warmed 48-well plate taking care to avoid air bubbles. For each medium, 1 mL of each media was prepared with five gemcitabine concentrations were prepared (0 nM, 10 nM, 100 nM, 1 μM, 10 μM). 250 μL of media was added to each well. Triplicate wells were prepared for each drug concentration in each medium. The 48-well plate was then incubated in a Sartorius/Essen BioScience IncuCyte S3 Instrument with the lid on and organoids were monitored using the Organoid QC Assay over four days. Organoid area from three randomly selected organoids per well was measured daily over 4 days using OrganoID software as previously described (9).

To test the impact of chemotherapeutics on patient-derived organoids derived in standard medium that were cultured in either TIFM or standard media, organoids in standard organoid medium were dissociated into single cells by dissolving the domes for 2.5 hours at 37 °C with Collagenase/Dispase (Roche, 10269638001) and afterwards digested into single cells using Accutase (Sigma-Aldrich, A6964) for 30 min on an orbital shaker. Cells were then seeded into white bottom 96-well plates pre-coated with Cultrex Reduced Growth Factor Basement Membrane Extract, Type 2, Pathclear (R&D Systems, 3533-001-02) diluted in 1:3 PBS (Gibco,

10010023), at a density of 2,000 cells per well in hCLPT medium or at 5,000 cells per well in TIFM. Twenty-four hours after seeding, cells were treated in triplicates with concentrations ranging from 0.013 to 50 µM of gemcitabine (Selleckchem, S1714). After 4 days of drug incubation, cell viability was analyzed using Cell Titer Glo 2.0 Assay (Promega, PRG9242) according to the manufacturer's protocol. Luminescence was measured for 1000 ms using a microplate reader (Varioskan LUX™, Thermo Scientific). Viability percentages were normalized to vehicle-treated cell viability.

#### *Clonetracr analysis*

The ClonTracer library (18) was obtained from Addgene (67267). 50,000 mPDAC3 cells in RPMI-1640 supplemented with 10% dialyzed FBS were barcoded by lentiviral infection at an MOI of 0.1 to ensure infection of 5,000 cells with one unique barcode each. Cells were then allowed to expand for 8 population doublings before selection with 2 µg/ml puromycin over the course of 3 days. After selection, cells were maintained at a coverage of 85 cells per guide. Barcode-expressing cells cultured in RPMI-1640 were transferred into RPMI (RPMI → RPMI) or gradually transitioned into TIFM (RPMI → TIFM) over a period of 17 population doublings. Cells in each condition were split at 90% confluency and refed every day with fresh media to avoid nutrient depletion. For transitioning cells into TIFM, cells were initially plated in 90% TIFM/10% RPMI until they reached confluency. Cells were then transitioned into 99% RPMI/1% TIFM until they reached confluency. Cells were then transferred into 100% TIFM and maintained under these conditions until 17 doublings were completed (PD17). PD17 for both conditions, as well as the day 0 control (PD0), were frozen for later analysis. Genomic DNA was isolated from frozen cell pellets using the NucleoSpin BloodXL kit (Machery-Nagel, 740950). PCR was used to amplify the barcode region along with introducing Illumina adapters with 5 bp-long index sequences to each sample as previously described (18). 1.6 µg of genomic DNA was used as input for each PCR reaction, with each sample having two reactions to ensure sufficient template coverage. PCR products were column purified (Qiagen, 28104) and sequenced on an Illumina NovaSeqX NGS sequencing platform at the University of Chicago Functional Genomics Facility (RRID:SCR_019196). Quality control and filtering by hamming distance to account for sequencing errors was performed as described previously (18). Briefly, FASTQ files were prepared and sequencing reads were filtered based on an average Phred quality score greater than 30. Barcodes differing by 1 base pair from a more abundant sequence occurring at less than 12.5% of its count, or 2 base pairs from a more abundant sequence occurring at less than 2.5% of its count, were reassigned to the more abundant barcode. Finally, only barcodes with at least two observations were included in the final dataset. Gini index was calculated using the ineq package in R. Histograms of barcode frequency were created using the top 2,500 barcodes in each condition. The relation in fitness between clones under RPMI → RPMI and RPMI → TIFM conditions was determined by initially calculating barcode abundance as the proportion of reads belonging to a unique barcode relative to the total reads within a sample. The $\log_2$ fold change in the abundance of each barcode between PD0 and PD17 was then calculated for both medium conditions. Lastly, Spearman's rank correlation was calculated between the change in abundance of a clone under RPMI → RPMI and RPMI → TIFM conditions.

#### *Alkaline comet assays*

Alkaline comet assays were performed using the Trevigen CometAssay kit (Bio-Techne 4250-050-K). mPDAC1 RPMI and TIFM cells were plated in 6-well tissue culture plates at a density of 200,000 cells per well and allowed to adhere overnight. The following day, cells were treated with chemotherapeutics (vehicle, 150 nM gemcitabine, 10 µM SN-38 or 5 µM 5-FU) and harvested after 18 hours of treatment. Adherent cells were washed once with PBS and detached using 1% Trypsin-EDTA (Gibco 15400054) in serum-free RPMI-1640 for 20 minutes at 37 °C. Resuspended cells are diluted in PBS, added to 37 °C molten low melt agarose, and spread over pre-coated CometAssay slides. Slides are placed in 4 °C to solidify the agarose for 30 minutes and placed in pre-chilled 4 °C CometAssay lysis buffer to lyse overnight. The following day, slides are placed in pre-chilled alkaline unwinding solution for 1 hour at 4 °C. Slides are then subjected to electrophoresis in chilled alkaline unwinding solution at 4 °C using 25 V for 35 minutes. Slides are rinsed twice with water for 5 minutes, once with 70% ethanol for 5 minutes and dried at 37 °C for 15 minutes. Slides are then stained with SYBR green (Thermo, S33102) diluted 1:10,000 in TE buffer (10 mM Tris-HCL pH 7.5, 1 mM EDTA) for 30 minutes. Slides are rinsed twice with water and stored at room temperature until analysis performed the following day. Widefield imaging of comet assay slides was performed using an Olympus IX81 inverted widefield microscope with a 20x objective capturing fluorescence signal with 480/40 nm excitation and 510 nm emission settings. A minimum of 100 cells per sample were imaged for each experiment. Images were analyzed in Fiji/ImageJ software using the OpenComet plugin (10). Comet finding was performed using background correction and the brightest region head finding approach. Valid comet finding was manually confirmed for each data point included in the analysis.

***Cell cycle analysis***

Cell cycle profiles were determined using the Click-iT EdU Flow cytometry kit (Thermo, C10424). Given the presence of unlabeled nucleosides in TIFM medium that may interfere with analysis of EdU incorporation, a minimal TIFM medium containing only the pools of metabolites comprising glucose, vitamins and amino acids was generated for use during the labeling period. mPDAC1 RPMI and TIFM cells were plated in 6-well tissue culture plates at a density of 200,000 cells per well and allowed to adhere overnight. The following day, relevant treatment conditions were applied and harvested at the following timepoints: 0 hr, 24 hr, and 48 hrs. Prior to sample collection, cells were rinsed once with PBS and pulsed with 10 µM EdU in RPMI or minimal TIFM medium for 45 minutes at 37 °C. Cells were then rinsed with PBS and detached using 1% Trypsin-EDTA (Gibco 15400054) in serum-free RPMI-1640 for 20 minutes at 37 °C. Samples were then fixed for 15 minutes at room temperature with Click-iT fixative reagent and permeabilized with the Click-iT saponin-based permeabilization and wash reagent for 15 minutes at room temperature. Click-iT reactions with Alexa Fluor 647 azide were performed for 30 minutes at room temperature and cells were then stained with 300nM DAPI (Thermo, D1306) for 15 minutes prior to analysis. Flow cytometry analysis was performed using a BD LSR Fortessa Instrument. Cells were gated based on the signal intensity of FSC-A and SSC-A channels. Singlets were then gated based on the intensity of FSC-H and SSC-A channels. Total DNA content was determined based on fluorescence intensity from the DAPI channel. Active DNA replication was determined based on fluorescence intensity from the AF647 channel. Cell cycle profiles of samples were determined using FlowJo analysis software. The population of cells in G1-phase was determined as the event population with low DAPI signal and low EdU signal, cells in G2/M-phase as the event population with high DAPI signal and low EdU signal, and cells in S-phase with intermediate DAPI and high EdU signal. Proportions were determined based on the sum of events from these 3 populations.

***Immunoblotting***

Cells were plated during log-phase of growth into 6-well plates at a density of 100,000 cells per well and allowed to adhere overnight. Cells were then washed with PBS and lysed on ice for 15 minutes using RIPA lysis and extraction buffer (Thermo, 89900) supplemented with Pierce protease inhibitor cocktail (Thermo A32963) and Pierce phosphatase inhibitor cocktail (Thermo, A32957). During drug treatment experiments that would result a loss of cell viability, floating cells in the medium were simultaneously isolated by centrifugation at 1000 x g for 5 minutes and added to the lysate of the adherent cells. Lysates were sonicated three times for 10 s using a probe sonicator. Insoluble fractions were then removed by centrifugation at 21,000 x g for 20 minutes at 4 °C. Protein concentration of lysates were determined by BCA assay (Thermo, 23225). Protein lysates (30–50 µg) were resolved on sodium dodecyl sulfate–poly-acrylamide gel electrophoresis (SDS-PAGE) 4–12% Bis-Tris Gels (Invitrogen, NP0321) by running at 85V for 2 hours and transferred to a polyvinylidene difluoride membrane using the iBlot 2 Dry Blotting System (Invitrogen, IB21001). Membranes were blocked with Intercept Blocking Buffer (LI-COR, 927-70001) at room temperature for 1 hour, and stained overnight at 4 °C with the following primary antibodies: BCL-XL (1:1000 dilution, Cell Signaling 2764S), BIM (1:750, Cell Signaling 2933S), BAX (1:750, Cell Signaling 14796S), BAK (1:750, Cell Signaling 12105), BCL2 (1:1000, Cell Signaling 3498), MCL1 (1:750, Cell Signaling 94296), Histone H2A.X (1:1000, Cell Signaling 7631S), phosho-H2A.X

(Ser139) (1:1000, Cell Signaling 9718), ACTB (1:2000, Proteintech 66009-1-Ig), cleaved CASP3 (Asp175) (1:1000, Cell Signaling 9661), COXIV (1:1000, Abcam, Ab16056), TUB1A (1:1000, Cell Signaling 3873T), MAPK3/MAPK1 (Erk1/2) (1:1000, Cell Signaling 4695), Phospho- MAPK3/MAPK1 (Erk1/2) (Thr202/Tyr204) (1:1000, Cell Signaling 9101). Membranes were washed 3 x 5 minutes with PBS-T buffer and stained with the following secondary antibodies for 1-2 hours at room temperature: IRDye 680LT Goat anti-Rabbit IgG (1:10,000 dilution, Licor 926-68020), IRDye 800CW Goat anti-Mouse IgG (1:10,000, Licor 926-32210). Fluorescent membranes were scanned using a Licor Odyssey CLX instrument. Approximate molecular weights of identified protein species were determined by comparison to migration of the Chameleon duo pre-stained protein ladder (Licor, 928-60000).

#### *Generation of Omi-mCherry expressing cell lines*

mPDAC1 cells were transduced with retroviral particles to express Omi-mCherry (Addgene, 48685) and transduced cells were selected by treatment with 2 µg/ml puromycin over the course of 3 days. Following puromycin selection, cells were sorted via using a BD FACSAria II cell sorter to ensure uniform expression of the Omi-mCherry reporter in the mPDAC1 cells.

#### *MOMP reporter assay*

Cells were plated in 12-well tissue culture plates at a density of 30,000 cells per well and allowed to attach overnight. Cells were then treated the following day with 125 nM Gemcitabine. After 48 hours, the supernatant was collected and centrifuged at 1000 x g for 5 minutes to pellet the floating cells. Simultaneously, cells still adherent to the plate were detached using 1% Trypsin-EDTA (Gibco, 15400054) in serum-free RPMI-1640 for 20 minutes at 37 °C. Detached cells in Trypsin-EDTA were combined with the floating cells and centrifuged at 1000 x g for 5 minutes. 300 nM DAPI (Thermo, D1306) was added to the resulting pellet and incubated on ice for 5 minutes to label dead cells. The cells were then washed once with PBS plus 1% BSA and resuspended in PBS. Cells were strained with 40 µm cell strainers and analyzed by flow cytometry on a BD LSR Fortessa instrument. A minimum of 10,000 events per sample were captured for data analysis conducted using FlowJo software. Cells were gated based on the signal intensity of FSC-A and SSC-A channels. Singlets were then gated based on the intensity of FSC-H and SSC-A channels. Viable cells were then gated based on negative DAPI labeling. Viable, post-MOMP cells were determined based on low red fluorescence from the Omi-mCherry reporter. Fluorescence intensity values from the Omi-mCherry reporter exhibited a bimodal distribution, and the threshold was chosen as the minimum between these two population peaks.

#### *Subcellular fractionation*

20,000,000 mPDAC1 TIFM and RPMI cells were plated in 15 cm dishes and allowed to adhere overnight. The following day, cells were treated with 250 nM gemcitabine and harvested at the following time points: 0 hr, 12 hr, and 24 hr. Subcellular fractionation was performed using the mitochondrial isolation kit for cultured cells (Thermo, 89874) using the dounce homogenization method. Protein concentrations from cytosolic and crude mitochondrial lysate fractions were measured by BCA assay (Thermo, 23225). Samples were then run on 4-12% Bis-Tris Gels and transferred as described above. Membranes were subjected to fluorescence-based immunoblotting using COXIV, BAX, and TUBA1A primary antibodies as described above.

#### *BH3 profiling assays*

We performed BH3 priming as described previously (11). Briefly, BH3 priming treatment plates were prepared in black 384-well plates containing 15 µL volume of MEB buffer (150mM D-Mannitol, 10mM HEPES pH 7.5, 50mM KCl, 5mM Succinate, 20µM EGTA, 20µM EDTA, 0.1% BSA) containing the indicated concentration of BIM peptide (Genscript), 2 µM JC-1 (Invitrogen, T3168), 100 µg/mL digitonin, 20 µg/mL oligomycin and 10 mM 2-mercaptoethanol. Detached mPDAC cell samples were resuspended in MEB buffer at 20,000 cells per 15 µL. 15 µL of mPDAC cells in MEB were transferred to each well in the 384-well plate for a total volume of 30

µL per well. The treatment plate was immediately loaded onto a pre-heated fluorescence plate reader (SpectraMax iD5 Multi-Mode Microplate Reader, Molecular Devices) set to 30 °C. Fluorescence of JC-1 was monitored using an excitation of 545nm and emission of 590nm. Relative fluorescence intensity was recorded every 5 minutes for 180 minutes. The percentage of mitochondrial membrane depolarization was determined based on the area under the curve (AUC) estimates for each sample well relative to the positive (FCCP, carbonyl cyanide-p-trifluoromethoxyphenylhydrazone) and negative (PUMA2A, a modified BH3-only peptide without MOMP-inducing activity obtained from Genscript) depolarization controls, using the following formula:
$$\%\,Depolarization = \left(1 - \frac{Sample_{AUC} - FCCP_{AUC}}{PUMA2A_{AUC} - FCCP_{AUC}}\right) * 100.$$

***Genome-wide CRISPR screening***

The genome-wide CRISPR–Cas9 screen was adapted from Colville et al. (19). Briefly, the CRISPR/Cas9 Brie sgRNA library was obtained as lentiviral suspension from Addgene (73633-LV). 250 million Cas9-expressing mPDAC1 TIFM cells were infected with the genome-wide library at an MOI of 0.3. Infected cells were then selected using puromycin (1ug/mL). Given that mPDAC-TIFM cultures maintain therapy resistance for ~1 week after a switch to RPMI medium, to facilitate lentiviral infection and the scale of the screen, mPDAC1-TIFM cells were maintained in RPMI plus 1mM glycine for the duration of the screen. 80 million infected cells were then plated and treated the following day with gemcitabine at 250nM. After 3 days of treatment, supernatants were recovered from the plates to collect the dead cell fraction. Plates were then washed with PBS, and trypsinized to collect the live cell fraction. Cell viability measurements for both fractions were collected using a Vi-Cell XR Counter and indicated a coverage of 875x and a 375x for the live and dead cell fractions, respectively. The dead and live fractions were then centrifuged for 10min at 550g and genomic DNA was harvested from cell pellets using the DNeasy Blood and Tissue Kit (Qiagen 69504). PCR amplification of sgRNA sequences from genomic DNA was performed as described in Doench et al. (20). PCR products from live and dead cell fractions were then submitted for next generation sequencing with the UChicago Genomics Core Facility and sequenced on an Element Biosciences AVITI platform. Data analysis of sequencing results was performed following the MAGeCK Counts pipeline (21) within the Galaxy web server (22). Relative abundances of individual sgRNAs within each sample were computed and the median was determined for each gene. Genes differentially required for survival under chemotherapy exposure were determined comparing sgRNA abundances between live and dead cell fraction using MaGeCK.

***CRISPR vector cloning and CRISPR-mediated gene knock out in cell lines***

CRISPR/Cas9 gene editing was utilized to generate knockout mPDAC cell lines for *Bcl2l1* and *Bax/Bak1*. The top 2 single guide RNAs (sgRNAs) from the Brie library (Doench et al., 2016) were selected to perturb *Bcl2l1*. Protospacer sequences targeting *Bcl2l1* or control non-targeting sequences purchased as single strand oligonucleotides from IDT using standard desalting purification. Oligonucleotides were then cloned into the lentiCRISPRv2-Blast vector (Addgene, 83480) using previously published methods (5). mPDAC1-TIFM cells were lentivirally transduced with the resulting vectors as previously described. Transduced cells were selected over the course of 5 days with treatment of 5 µg/ml blasticidin. Following selection, *Bcl2l1* knockout was confirmed by immunoblotting for BCL-XL. BCL-XL knockout proved highly efficient in mPDAC1 cells and therefore cell viability and immunoblotting assays were performed using this polyclonal cell population.

Dual targeting of *Bax/Bak1* was accomplished by cloning two small guide RNA and promoters into the lentiCRISPRv2 vector (Addgene, 52961) as previously described (6). The top sgRNAs against *Bax* and *Bak1* were selected from the Brie library (Doench et al., 2016) and utilized to design dsDNA fragments (IDT) containing the first sgRNA (whose expression is driven by constitutive expression from the U6 promoter contained in the lentiCRISPRv2 vector backbone), H1 promoter (to drive constitutive expression of a second sgRNA) and the second sgRNA. The dsDNA fragment was flanked by BsmBI restriction sites to facilitate subcloning into lentiCRISPRv2. dsDNA fragments were then cloned into lentiCRISPRv2 as described above. mPDAC1-RPMI cells were lentivirally transduced with the above dual-targeting vectors (or control vectors containing two non-targeting sgRNAs). Single sgRNA knockout of *Bax/Bak1* proved inefficient in mPDAC1 cells. Work from other groups suggests that targeting two sgRNAs against a single gene can improve efficiency in CRISPR systems (7). Therefore, we cloned additional *Bax/Bak1* dual targeting vectors into the lentiCRISPRv2-Blast vector (Addgene, 83480). We then transformed mPDAC1 cells with both lentiCRISPRv2-Blast and lentiCRISPRv2 vectors and selected cells using both blasticidin (5 μg/ml) and puromycin (2 μg/ml) over the course of 5 days. *Bax* and *Bak1* knockout was confirmed by immunoblotting and we observed high knockout efficiency using this approach. Cell viability assays were performed using this polyclonal cell population.

All sgRNA sequences used in this study are listed in the table below:

| Gene ID | sgRNA | Protospacer Sequence |
|---|---|---|
| *Bcl2l1* | 1 | AGTAAACTGGGGTCGCATCG |
| *Bcl2l1* | 2 | CTGCTCAAAGCTCTGATACG |
| *Bax* | 1 | GACACGGACTCCCCCCGAGG |
| *Bax* | 2 | CTGATGGCAACTTCAACTGG |
| *Bak1* | 1 | GGTAGACGTACAGGGCCAGA |
| *Bak1* | 2 | GGAACTCTGTGTCGTAGCGC |
| *YFP* | 1 | CAACTACAAGACCCGCGCCG |

***Co-immunoprecipitation assays***

For co-immunoprecipitation assays, 1,000,000 mPDAC1 TIFM and RPMI cells were seeded per condition in 10cm plates and allowed to adhere overnight. The following day, cells were treated with vehicle (DMSO), 125 nM Gemcitabine, or 125 nM Gemcitabine plus 100 nM A-1331852. After 24 hours of treatment, media was collected and floating cells were pelleted by centrifuging at 1000 x g for 5 minutes. Adherent cells were washed with 2 mL PBS and lysed using complete Co-IP lysis buffer (0.2% NP-40, 50 mM Tris-HCL, 150 mM NaCl, 1 mM EDTA, and 10% glycerol at pH 7.5 and supplemented with Pierce protease inhibitor cocktail (Thermo, A32955) and Pierce phosphatase inhibitor cocktail (Thermo, A32957)) and incubated on ice for 15 minutes. Lysates from adherent and floating fractions were combined and the insoluble fraction was removed by centrifugation at 21,000 x g for 20 minutes at 4°C. Protein concentrations of lysate were determined by BCA assay (Thermo, 23225) and 1 mg total protein lysate was used as input for co-immunoprecipitation reactions. Samples were initially pre-cleared with Protein A/G Agarose beads (Santa Cruz, sc-2003), rotating for 1hr at 4 °C. 1 μg of BCL-XL IgG antibody (Cell Signaling 2764) or 1 μg of Rabbit IgG Isotype Control (Cell Signaling 3900) was added to lysates, which were then rotated overnight at 4°C.

The following day, 20 μl of Protein A/G Agarose beads were added to lysates to capture antibody:protein complexes and rotated for 3 hours at 4°C. Agarose beads were pelleted by centrifugation at 1000 x g for 3 minutes at 4 °C and the supernatant removed. Samples were washed 3x with complete Co-IP lysis buffer and eluted using complete Co-IP lysis buffer plus 1x SDS-PAGE sample loading buffer (Thermo J61337AD) by boiling samples at 100 °C for 5 minutes. Samples were run on 4-12% Bis-Tris Gels and transferred as described above. Membranes were blocked with Intercept Blocking Buffer (LI-COR, 927-70001) at room temperature for one hour and incubated overnight with the primary antibodies listed above. Membranes were then washed 3 x for 5 minutes with PBS-T and then incubated with Mouse Anti-rabbit Conformation Specific IgG (1:2000 dilution, Cell Signaling 3678) for one hour at room temperature. Membranes were washed 3 x for 5 minutes with PBS-T and then incubated with Anti-mouse IgG, HRP-linked Antibody (1:1000 dilution, Cell Signaling 7076) for one hour at room temperature. Membranes were treated with SuperSignal™ West Dura reagent (Thermofisher, 34075) for 2 minutes and then chemiluminescence was imaged using an Invitrogen iBright FL1000 digital imager. The molecular weights of protein bands were determined based on migration relative to visible marker bands in Chameleon duo pre-stained protein ladder (Licor, 928-60000).

*Generation of mPDAC-RPMI→TIFM and mPDAC-RPMI→low arginine cultures*

To generate low arginine medium, RPMI-1640 without amino acids and glucose (US Biological Life Sciences, D9800-27) was supplemented with all amino acids and glucose to RPMI-1640 concentrations, except for arginine which was supplemented at a concentration of 10 µM. Murine cancer cell lines described above were then cultured in this low arginine RPMI media for 6 weeks prior to the start of experiments.

To generate mPDAC-RPMI→TIFM cultures, murine and human PDAC cell lines were initially transferred to a medium environment containing 99% TIFM and 1% RPMI for about 2 weeks until cultures grew to confluency. These cultures were then passaged in 100% TIFM. Cells were then maintained in TIFM for 4 weeks prior to that start of experiments.

*Tumor adapted culture generation*

Orthotopic allograft tumors were established with mPDAC3-RPMI cells as described above. 4-6 weeks after implantation, tumors were processed for cancer cell isolation. 4 tumors were pooled as a single sample and dissociated as described previously (1). Single cell tumor suspensions were plated onto 10 cm tissue culture plates in RPMI-1640 medium with 10% dialyzed FBS supplemented with penicillin-streptomycin (ThermoFisher, 15140-148) and gentamicin (ThermoFisher, 15750060). The tumor cell suspension was then allowed to adhere and acclimate to standard culture conditions for 48 hours. To isolate cancer cells from the total tumor cell population, tumor cells were detached using 1% Trypsin-EDTA (Gibco, 15400054) in serum-free RPMI-1640 for 20 minutes at 37°C. Samples were then blocked using 10% normal mouse serum (Thermo, 10410) in PBS containing 1% BSA. Samples were then stained with rat anti-mesothelin (mouse) (MBL, D233-3) antibody at 1:100 dilution in 10% normal mouse serum in PBS containing 1% BSA for 30 minutes at 4 °C. Samples were washed twice with PBS containing 1% BSA and then stained with APC-conjugated goat anti-rat IgG (Thermo, A10540) secondary antibody at 1:200 dilution in 10% normal mouse serum in PBS containing 1% BSA for 20 minutes at 4 °C. APC-positive cancer cells were sorted via FACS using a BD FACSAria II Cell Sorter and returned to standard culture conditions supplemented with penicillin-streptomycin and gentamicin pen/strep and gentamicin for 48 hours. After 48hrs, sorted tumor-adapted cells were replated for subsequent analyses.

*Propidium iodide-exclusion viability assay*

Cell viability was assessed in certain experiments by propidium iodide staining coupled with flow cytometry, which also determines cell viability based on loss of plasma membrane integrity. mPDAC cells were plated in 6-well tissue culture dishes at a density of 100,000 cells per well and allowed to adhere overnight. The following day, the medium on each culture was changed and included compounds at indicated concentrations of vehicle. After indicated periods of drug treatment, propidium iodide (thermos P3566) was added to the culture medium at a concentration of 1.5 µg/ml and cultures were incubated at 37°C for 30 minutes to label dead cells. Afterwards, the medium was collected, and the dead cell fraction was isolated by centrifugation at 1000 x g for 5 minutes at 4 °C. Simultaneously, adherent cells were detached using 1% Trypsin-EDTA (Gibco 15400054) in serum-free RPMI-1640 for 20 minutes at 37 °C. The detached cell and floating cell fractions were combined and washed 3 times with PBS containing 1% BSA. Cells were strained with 40 µm cell strainers and analyzed by flow cytometry on a BD LSR Fortessa instrument. A minimum of 10,000 events per sample were captured for data analysis conducted using FlowJo software. Cells were gated based on the signal intensity of FSC-A and SSC-A channels. Singlets were then gated based on the intensity of FSC-H and SSC-A channels. Viable cells were then gated based on negative propidium iodide labeling, and thresholding for positive staining was determined based on the fluorescence intensity of an unlabeled cell sample

*Drugs*

*In vitro* cell viability assays were performed using the following compounds: Gemcitabine hydrochloride (Sigma, G6523), SN-38 (Cayman, 15632), 5-Fluorouracil (Cayman, 14416), Palbociclib (Cayman, 16273), Paclitaxel (Cayman, 10461), Oxaliplatin (Cayman, 13106), Trametinib (Cayman, 16292), RMC-7977 (MedChem Express, HY-156498), S63845 (Selleck Chem, S8383), Venetoclax (Selleck Chem, S8048), Navitoclax (Selleck Chem, S1001), A-1331852 (Selleck Chem, S7801). *In vivo* studies were performed using the following compounds: Gemcitabine hydrochloride (USP, 1288463).

***Lentivirus and retrovirus production***

Lentiviral transduction was used to generate CRISPR-mediated knockout cell lines and Nuclight expressing cell lines. Lentivirus was produced using 3$^{rd}$ generation packaging systems. HEK293T cells cultured in DMEM/F12 medium with 10% FBS were transfected with 6 μg of the relevant transfer plasmid, 1.5 μg of pMD2.G (Addgene, 12259), 3 μg of pMDLg/pRRE (Addgene, 12251), and 1.5 μg of pRSV-REV (Addgene, 12253) using TransIT-LT1 transfection reagent (MirusBio, MIR 2304). Media was replaced after overnight transfection and harvested for viral particle collection after 48 hours. Collected virus was filtered though 45μm sterile syringe filters and stored at -80°C time of lentiviral transduction.

Ecotropic retrovirus was utilized for the generation of Omi-mCherry expressing cell lines. Retrovirus was generated by transfecting HEK293T with 5 μg the pBabe-Omi-mCherry transfer plasmid (Addgene, 48685) and 5 μg of pCL-Eco packaging plasmid (Addgene, 12371) using TransIT-LT1 transfection reagent (MirusBio, MIR 2304). Media was replaced after overnight transfection and harvested for viral particle collection after 48 hours. Collected virus was filtered though 45μm sterile syringe filters and stored at -80°C time of lentiviral transduction.

**Sex as a biological variable**

Our study was carried out in female mice. While pancreatic cancer affects males and females with similar frequency, we made use of female mice as the pancreatic cancer cell line we use for allograft animal experiments was derived from a female mouse.
